# Single-cell epigenomic reconstruction of developmental trajectories in human neural organoid systems from pluripotency

**DOI:** 10.1101/2023.09.12.557341

**Authors:** Fides Zenk, Jonas Simon Fleck, Sophie Martina Johanna Jansen, Bijan Kashanian, Benedikt Eisinger, Małgorzata Santel, Jean Dupre, J. Gray Camp, Barbara Treutlein

## Abstract

Human cell type diversity emerges through a highly regulated series of fate restrictions from pluripotent progenitors. Fate restriction is orchestrated in part through epigenetic modifications at genes and regulatory elements, however it has been difficult to study these mechanisms in humans. Here, we use organoid models of the human central nervous system and establish single-cell profiling of histone modifications (H3K27ac, H3K27me3, H3K4me3) in organoid cells over a time course to reconstruct epigenomic trajectories governing cell identity acquisition from human pluripotency. We capture transitions from pluripotency through neuroepithelium, to retinal and brain region specification, as well as differentiation from progenitors to neuronal and glial terminal states. We find that switching of repressive and activating epigenetic modifications can precede and predict decisions at each stage, providing a temporal census of gene regulatory elements and transcription factors that we characterize in a gene regulatory network underlying human cerebral fate acquisition. We use transcriptome and chromatin accessibility measurements in the same cell from a human developing brain to validate this regulatory mode in a primary tissue. We show that abolishing histone 3 lysine 27 trimethylation (H3K27me3) through inhibition of the polycomb group protein Embryonic Ectoderm Development (EED) at the neuroectoderm stage disrupts fate restriction and leads to aberrant cell fate acquisition, ultimately influencing cell type composition in brain organoids. Altogether, our single-cell genome wide map of histone modifications during human neural organoid development serves as a blueprint (https://episcape.ethz.ch) to explore human cell fate decisions in normal physiology and in neurodevelopmental disorders. More broadly, this approach can be used to study human epigenomic trajectory mechanisms in any human organoid system.

**Summary:** Unguided neural organoids reveal widespread and dynamic switching of epigenetic modifications during development and recapitulate fate restriction from pluripotency to terminally differentiated cells of the human central nervous system.

---

During development and differentiation, cells undergo remarkable cell fate transitions starting from pluripotent cells and multipotent progenitors to give rise to all major cell types of the body. During these transitions, developmental plasticity becomes increasingly restricted until the cells stop dividing and acquire their terminal fate. Despite recent progress in the field of epigenetics and reprogramming, the details for how human cells acquire and maintain their identity remains elusive. What is known is that during differentiation, histone modifications are involved in the packaging of DNA on nucleosomes and can either directly or indirectly influence the expression of underlying genes^1^. *In vivo*, chromatin regulation during differentiation is dynamic and complex and recent studies have shown that mutations within histone tails or enzymes that modify the histone tails (including so-called readers, writers and erasers of histone modifications) are involved in neurodevelopmental disorders, which emphasize their importance in the developing nervous system^2,3^. Previous bulk studies of histone modifications in human developing tissues failed to capture the individual trajectories of furcating fates^4^. Here, we explore how epigenetic changes on chromatin are involved in cell fate decisions during the human brain and retina development. We model central nervous system development using organoids, and select three histone modifications as proxies for dynamic epigenetic change and validate our findings in a primary developing human brain. We include H3K27me3 as a repressive mark at developmental genes^5^, H3K27ac a mark of active enhancers and promoters^6,7^, and H3K4me3 as a mark of active or bivalent promoters of developmental genes^5,8,9^. In addition, these marks interact with one another and can act in concert or be mutually exclusive^10,11^. We provide an atlas of these marks over organoid development, identify different modes of epigenetic regulation during fate restriction and neuronal specification. We integrate all modalities and explore dynamics over differentiation trajectories from pluripotency using gene regulatory network analysis, and find that perturbing the writing of one of these marks (H3K27me3) in the early neuroepithelium results in loss of fate restriction and emergence of aberrant cell states.

## Results

### Single-cell epigenomic atlas of brain organoid development

To dissect the role and dynamic turnover of histone modifications during human brain and retinal organoid development, we performed CUT&Tag and mRNA-sequencing in single cells (scCUT&Tag and scRNA-seq) on a time course, covering cell fate transitions from iPSC to terminally differentiated neurons and glial cells. We recorded three histone modifications (H3K27ac, H3K27me3, H3K4me3) and RNA expression from the same cell suspensions at six developmental time points in brain organoids (day 5, 15, 35, 60, 120, 240) and two time points in retina organoids (day 45, 85) (Fig. 1a and Extended Data Fig. 1a and b). Retinal cells emerge from the developing diencephalic neuroepithelium and can spontaneously arise in un-guided brain organoids, supporting the inclusion of both organoid systems in a neural epigenomic trajectory reconstruction.

**Fig. 1.**
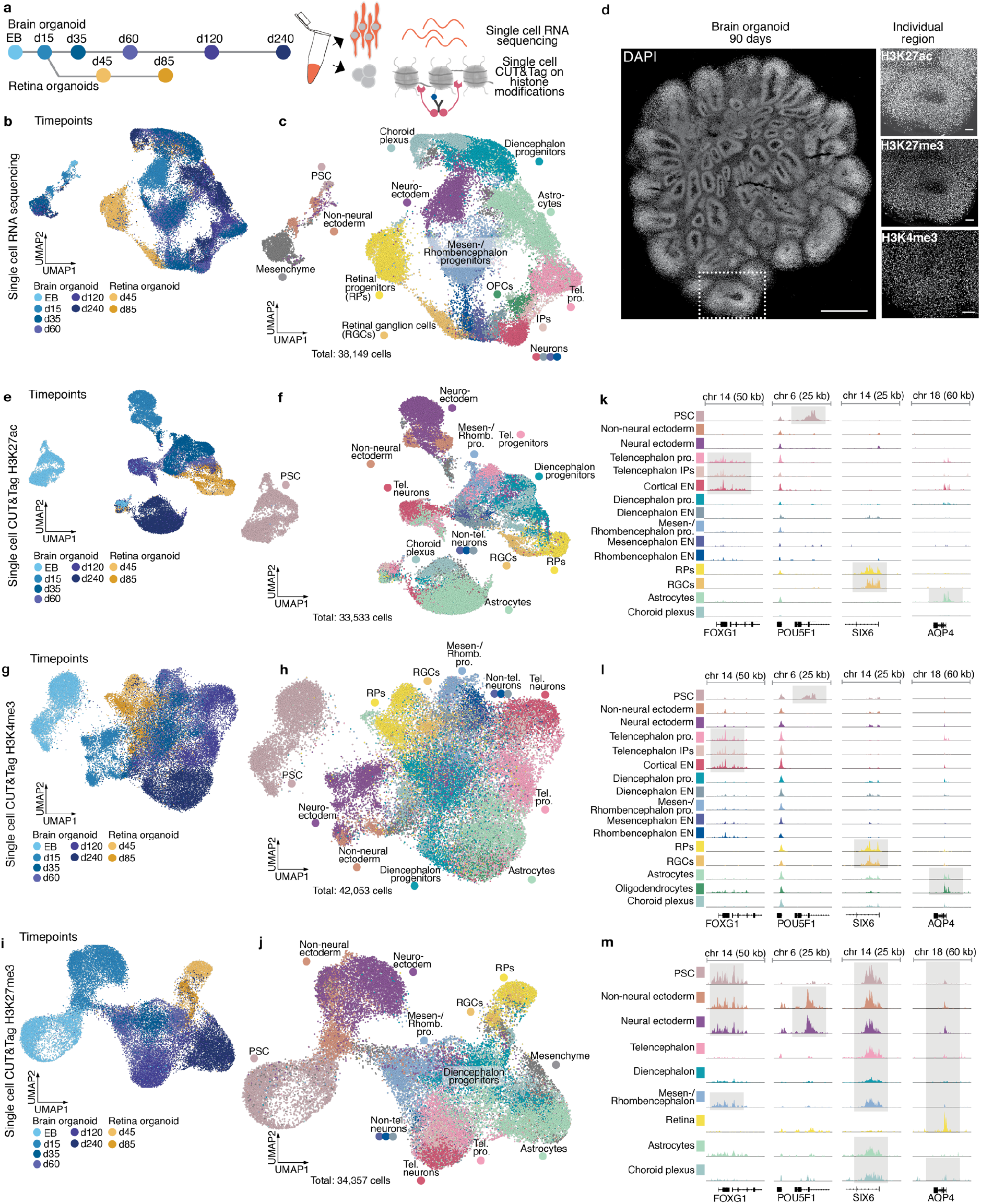
Single-cell epigenomic atlas of human brain organoid development from pluripotency to neurogenesis. a, Experimental outline. scCUT&Tag and scRNA-Seq were performed at different developmental time points during brain and retina organoid development. b, UMAP embedding of scRNA-seq data with cells colored based on developmental time point. c, UMAP embedding of scRNA-seq data with cells colored and labeled by cell state (IP - intermediate progenitor). d, DAPI staining of an organoid at day 90, scale bar 1000 µm (left panel). One ventricle of an organoid stained for H3K27ac, H3K27me3, H3K4me3, scale bar 100 µm (right panel) (Methods). e-j, UMAP embedding of single-cell CUT&Tag data for H3K27ac (e,f), H3K4me3 (g,h), and H3K27me3 (i,j) colored and labeled by time point (e, g, i) or cell state (f, h, j). k-m, Genome browser snapshots of the enrichment of the respective mark at four different marker genes (FOXG1-telencephalon, POU5F1-pluripotency, SIX6-retina, AQP4-astrocytes). Each signal track represents the summarized signal of all cells of the annotated cell state.

Dimensionality reduction and embedding with Uniform Manifold Approximation and Projection (UMAP)^12^ using the scRNA-seq data (38,149 cells) revealed diverse populations at each time point. We annotated clusters by comparing gene expression to reference datasets^13,14^ and analyzing marker gene expression. The cells represented in the dataset cover transitions from early pluripotent stages at d5 to a stratified neuroepithelium at d15, with progenitors diversifying into retina and brain regional identities (telencephalon, diencephalon, non-telencephalon) between days 35-60 (Fig. 1b and c). Brain region-specific and retinal neurons start to develop from day 35, increase in abundance over time (Fig. 1b and c), and by day 120 astrocytes and oligodendrocyte precursor cells (OPCs) appear, coinciding with the gliogenic switch during the second trimester of human embryonic development (Fig. 1b and c)^15,16^.

We investigated the spatial distribution of H3K27ac, H3K27me3, and H3K4me3 by immunofluorescence and found that all nuclei stain for the three histone modifications (Fig. 1d). We established scCUT&Tag in brain organoids (see Methods) and recorded the genome-wide distribution of H3K27ac (33,533 cells), H3K27me3 (34,357 cells), and H3K4me3 (42,053 cells) in the same cell suspensions as used for the scRNA-seq experiments. We obtained high-quality data from all experiments as reflected by the nucleosome pattern and the comparably high average number of fragments (H3K4me3, 2857; H3K27ac, 1723; H3K27me3, 784) recovered from each cell (Extended Data Fig. 1c and d)^17–19^. Dimensionality reduction and embedding with UMAP revealed remarkable cell state diversity over the time course for each modality (Fig. 1e-j).

To annotate cell states within the epigenomic datasets, we first performed a high-resolution Louvain clustering^20^ for each modality separately in order to enhance robustness through increasing fragment counts per modality (H3K27me3, 104k mean fragments; H3K27ac, 236k; H3K4me3, 417k). Next, we compared each epigenomic high-resolution cluster with the annotated scRNA-seq clusters using minimum-cost, maximum-flow (MCMF) bipartite matching^21^ based on correlation for activating histone marks (H3K27ac and H3K4me3) and anti-correlation for the repressive mark H3K27me3 (Extended Data Fig. 2a and b). Based on these matches we transferred annotated cell labels from RNA to the other modalities. After label transfer, cell state proportions were similar between all modalities, supporting integration accuracy (Extended Data Fig. 2c). In addition, we detected highly enriched signal (peaks) of activating marks at known regulators of cell type and regional identity (e.g. FOXG1, telencephalic branch; NEUROD2, telencephalic neurons; SIX6, retinal branch; LHX5, neuroepithelium and non-telencephalon neurons; HOXB2, rhombencephalic branch; POU5F1, pluripotent stem cells; AQP4, astrocytes) (Fig. 1k and l, Extended Data Fig. 2d) and found H3K27me3 signal (repressive) in cell states that lack expression of the respective genes (Fig. 1m). These data provide the first comprehensive epigenomic atlas of human central nervous system development in organoids containing activating and repressive histone modifications along with RNA expression on the single cell level. This atlas can also be browsed interactively at: https://episcape.ethz.ch.

### Epigenomic switches during regional diversification

To visualize the branching of neuroepithelial cells into neuronal progenitor cells (NPCs) and neurons of different brain regions, we computed terminal fate probabilities for each regional identity using CellRank^22^ based on RNA expression. We summarized these probabilities for each high-resolution cluster and used them to construct a graph representation of the differentiation events (Fig. 2a). A force-directed layout of this graph revealed the transition of pluripotent stem cells (PSCs) into neuroepithelium (NE), which then diversified into branches of region-specific neurogenesis (telencephalon, Tel; diencephalon, Dien; mesen-/rhombencephalon, MRh: retina, Ret) (Fig. 2b). By mapping the matched high-resolution clusters of chromatin modalities (Fig. 2a and Extended Data Fig. 2b) onto this representation, we could visualize histone modifications switching between neuroepithelium and brain region branches (Fig. 2c). For each chromatin modification, we computed a gene activity score by summarizing fragment counts over the gene body plus an extended promoter region. Visualizing gene activities on the graph layout revealed that region-specific transcription factors (e.g. FOXG1, NEUROD2, LHX5, VSX2) tend to be broadly repressed by H3K27me3 outside of their expression domain and show low levels of H3K4me3 (Fig. 2c). Overall, we found that most protein coding genes were either expressed or repressed at some point in the developmental time course (61%), some were never repressed and always active (16%), always inactive and repressed (15%) or not repressed and inactive (8%). Inactive genes were enriched for GO-terms for sensory perception, immune system as well as skin development and fertilization, indicating repression of non-neural programs.

**Fig. 2.**
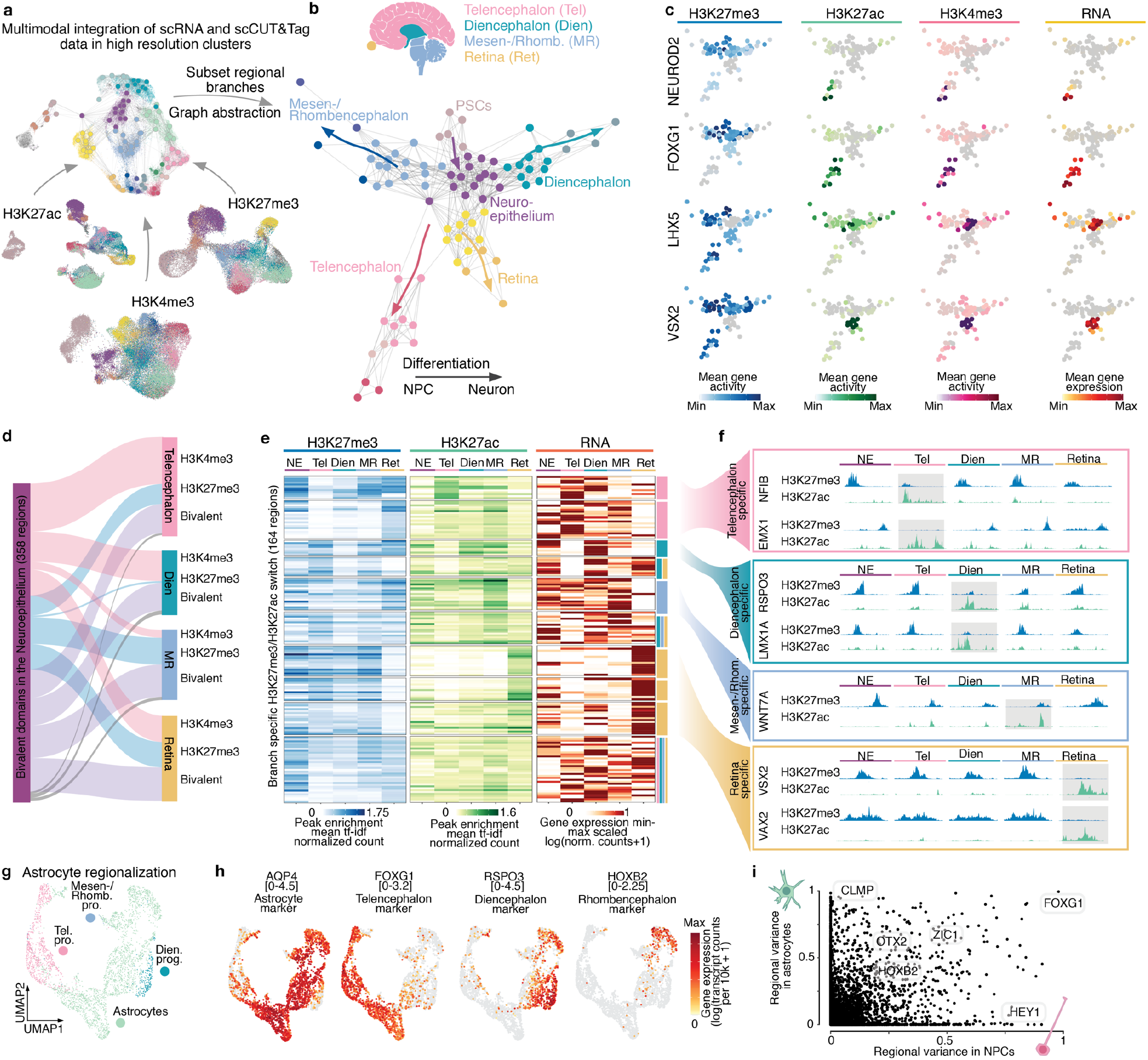
Integration of chromatin and RNA modalities reveals epigenetic switches during brain region diversification. a, Schematic of the multimodal data integration through high-resolution cluster matching between histone marks and RNA. b, Schematic of the human brain (top) and force directed-layout (bottom) of the matched high-resolution clusters colored by regional identities and cell state. c, Force directed-layout colored by RNA expression and gene activities of all three chromatin marks for NEUROD2, FOXG1 (telencephalon), LHX5 (Non-telencephalon) and VSX2 (retina). d, Alluvial plot of bivalent H3K27me3 and H3K4me3 peaks at the neuroepithelium stage. Most peaks get resolved and become enriched for either of the two marks (blue-H3K27me3 and pink-H3K4me3). Peaks that stay bivalent in any of the lineages are labeled in purple. e, Heatmap of signal enrichment of lineage-specific peaks showing switching between H3K27me3 and H3K27ac. Expression of the closest gene is shown in the right panel. f, Genomic tracks showing switching peaks close to lineage-specific genes (blue-H3K27me3, green-H3K27ac). g, UMAP embedding of NPCs and astrocytes from the 120 and 240 day time points within the dataset. h, UMAP embedding colored by gene expression of a general astrocyte marker (AQP4) and regional marker genes (FOXG1 - telencephalon), (RSPO3 - diencephalon), (HOXB2 - rhombencephalon). i, Scatter plot showing the regional variance of gene expression for NPCs and astrocytes.

Next, we sought to understand global chromatin modification changes as brain regions emerge from the neuroepithelium. We identified histone modifications (peaks) differential between all regional and cell type branches (Extended Data Fig. 3a-c) (Supplementary Table 1). Ontology analysis^23^ of genes in close proximity to H3K4me3 and H3K27ac peaks revealed a strong enrichment in developmental processes. In particular, activating H3K4me3 and H3K27ac peaks had enrichments for receptor signaling and metabolic processes (PSCs), embryonic development and morphogenesis (NNE - non-neural ectoderm), and patterning and nervous system development (NE). Within the regional branches, activating peaks were found enriched near genes associated with neuron maturation and differentiation (Tel), brain regional development (MRh), eye and sensory organ development (Ret), gliogenesis and glia cell differentiation (Ast - astrocytes), cerebrospinal fluid production as well as metabolic and immune processes (Chp - choroid plexus) (Extended Data Fig. 3a and b). In contrast to H3K4me3 and H3K27ac modifications, H3K27me3 domains repressed developmental regulators not involved in brain development. We found H3K27me3 peaks close to genes associated with heterochromatin assembly or regulation of cellular stress (PSC), mesenchymal stem cell proliferation (NNE), BMP signaling (Tel) and regulation of other cell fates such as pancreatic cells or keratinocytes (Dien and MRh) (Extended Data Fig. 3c) (Supplementary Table 2).

We observed abundant bivalent domains within the neuroepithelium, which are defined by H3K27me3 and H3K4me3 co-enrichment at repressed genes and have been reported to preferentially occur at genes that require rapid activation during development^8,9,24^ (Fig. 2c and d and Extended Data Fig. 3d-h). Bivalent domains became activated (installation of H3K4me3, removal of H3K27me3) predominantly in the telencephalon and diencephalon branch, while the majority of bivalent peaks acquired repression (H3K27me3) in the mesen-/rhombencephalon (Fig. 2d). A smaller subset of domains remained bivalent as cells transition into the regional branches. We next analyzed switching between H3K27me3 and H3K27ac, which are considered mutually exclusive as they are installed on the same histone H3 lysine 27 residue and rarely co-occur within a nucleosome^5,6,11^. We identified a set of peaks that were repressed (H3K27me3) during the neuroepithelial stage and switched into an active state (depletion of H3K27me3, enrichment of H3K27ac) in individual regional branches (Fig. 2e). This epigenetic activation was accompanied by region-specific expression of nearby genes (Fig. 2e), including important regulators of regional identity, such as EMX1 and NFIB (Tel), RSPO3 and LMX1A (Dien), WNT7A (MRh) and VSX2 and VAX2 (Ret) (Fig. 2f). We found that 60% of these genes showed previous bivalency within the neuroepithelium (Supplementary Table 3-4). We generated single cell multiome (linked scRNA-expression and scATAC-chromatin accessibility) and bulk CUT&Tag data from a primary developing forebrain at 19 gestational weeks (gw) (Extended Data Figure 4a-d). We found the same enrichment of activating and repressing histone modifications on these regional regulators (Extended Data Fig. 4d).

Interestingly, we observed region-specific gene expression in astrocytes within the organoids at 4 and 8 months (Fig. 2g and h). We calculated the regional variance of gene expression between NPCs and astrocytes from different regions (Fig. 2i) and found that FOXG1, OTX2, HOXB2, LHX2 as well as the ZIC- and IRX family define regionalization in both neuron and astrocyte populations, similar to observations in mouse and primary human tissue^25–29^ (Extended Data Fig. 5a-g). This is also reflected on the chromatin level when we associate region-specific peaks of the active marks H3K27ac and H3K4me3 to the closest gene (Extended Data Fig. 5h). Interestingly, H3K27me3 at regional marker genes (LHX1, FOXG1, LHX2) showed higher gene repression variance in NPCs compared to astrocytes indicating that gene repression in astrocytes is less predictive of regional identity. Altogether, our analysis reveals that the interplay of chromatin modifications induces and stabilizes brain region diversification events.

### Epigenetic activation precedes gene expression during neurogenesis

We next explored histone modification dynamics during differentiation from pluripotent stem cells to region-specific neurons. Focusing on the dorsal telencephalon trajectory, we ordered cells along pseudotime using RNA velocity^22^ for RNA, and diffusion maps^30^ for histone marks. To obtain an even time point distribution over pseudotime for all modalities, we sub-sampled cells before grouping them into 50 equally sized bins.

First, we used the pseudotime reconstructions to gain broad insight into chromatin domain changes throughout the development of the dorsal telencephalon. As expected, we found that pluripotent cells show enrichment of bivalent chromatin domains^8^ (Extended Data Fig. 6a), which decrease along with a gain of repressed (H3K27me3) and active (H3K27ac) domains until the NPC stage. Interestingly, bivalent domains increase in abundance again at the end of the trajectory when NPCs differentiate into neurons (Extended Data Fig. 6a). We validated the detected signal with our bulk CUT&Tag data from the human developing cortex (Extended Data Fig. 4a-d and 6b-d) and found that the chromatin status of domain groups was consistent with the organoid data.

Next, we clustered genes based on their pseudotemporal expression and histone modification patterns and found six major gene groups with distinct pseudotemporal dynamics (Fig. 3a., Supplementary Table 5 contains the full list of clusters). Clusters 1 and 2 (GO enrichment: structure formation, cell adhesion, and signaling receptor activity) capture the epigenetic silencing of pluripotency genes during the transition from pluripotency to neuroepithelium (Fig. 3a) following two different pseudotemporal dynamics: Genes in cluster 1 become downregulated and lose active histone modifications while gaining H3K27me3 at the neuroepithelium stage (Fig. 3b; POU5F1), whereas genes in cluster 2 install H3K27me3 only at the NPC stage while expression decreases right after exiting pluripotency (Fig. 3b; FOXO1). Cluster 3 genes (Fig. 3a, GO enrichment: epithelium, and central nervous system development) are expressed in the neuroepithelium and in neural progenitors and show transcription reduction concurrently with gain in H3K27me3 (Fig. 3b; LHX5). In contrast, genes in clusters 4 and 5 (Fig. 3a, GO enrichment: regionalization, telencephalon development, cerebral cortex development) are also expressed in NPCs, but showed an increase in active histone modifications and expression after H3K27me3 levels decreased (Fig. 3b; POU3F3). Strikingly, we observed that neuronal genes in cluster 6 lose H3K27me3 mediated repression and accumulate H3K27ac and H3K4me3 marks before RNA expression initiates (Fig. 3b-f; NEUROD2, GRIA2). These results support a model in which chromatin can be primed with activating histone modifications directing the future expression of target genes^1,31^. To consolidate this analysis and ameliorate the influence of potential confounders like distribution of time points and data quality, we repeated the pseudotime inference for the single time point day 120 after strict quality filtering using diffusion maps for all modalities (Extended Data Fig. 7a-d). After clustering we obtained again a cluster enriched in neuronal genes (Extended Data Fig. 7b, cluster 5, Supplementary Table 6), that showed priming with activating marks prior to RNA expression.

**Fig. 3.**
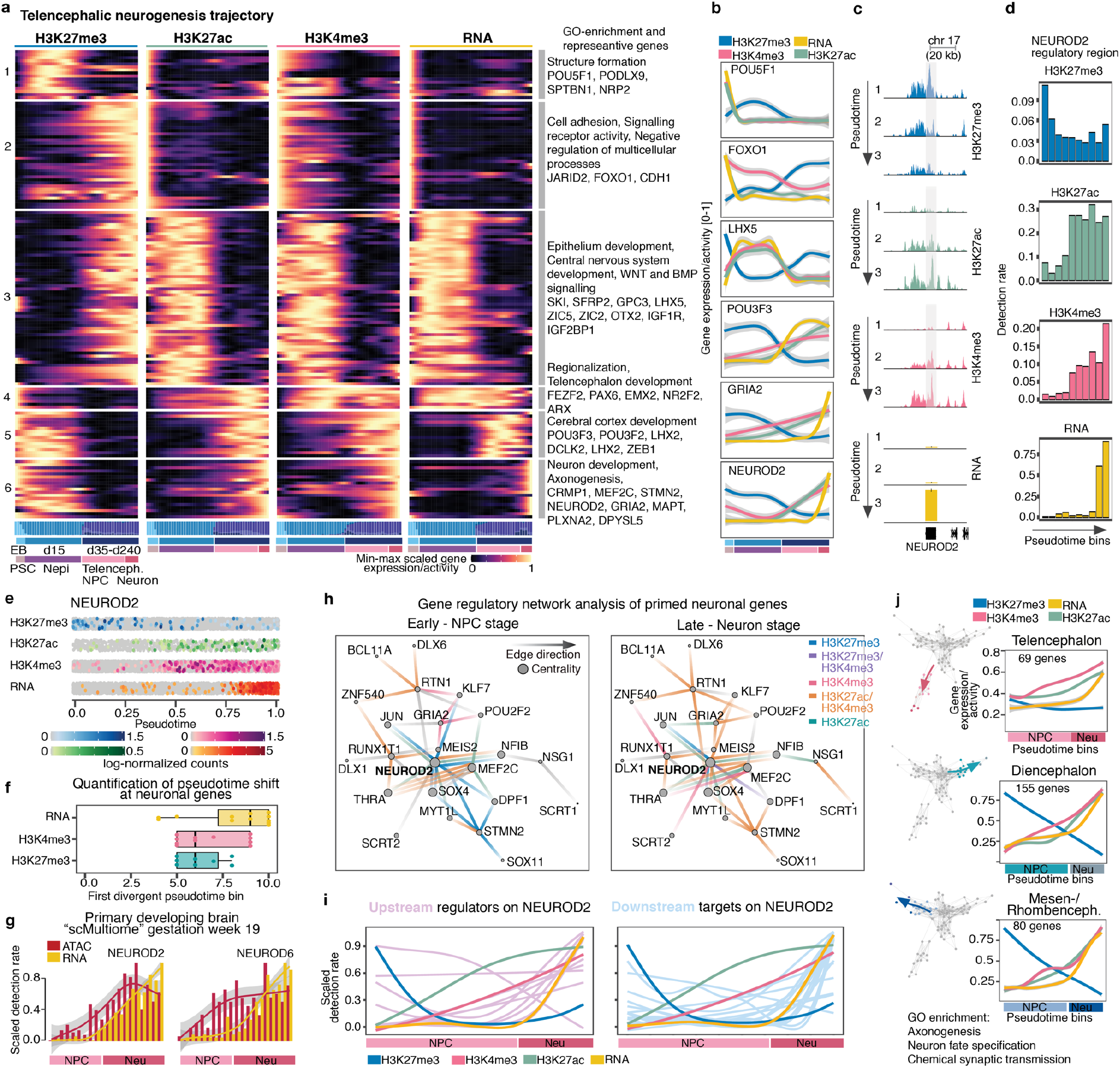
Pseudotime reconstruction from pluripotency reveals epigenetic priming during neuronal differentiation. a, Heatmap (left) showing gene activity scores for H3K27me3, H3K27ac and H3K4me3 as well as RNA expression over the telencephalic neuron differentiation trajectory from pluripotency. Pseudotime was binned and genes were K-means clustered based on average expression/activity of all marks and RNA in all bins. All clusters are annotated with representative GO-terms and example genes (right). Bar plots below show the distribution of the different time points (Supplementary Table 5 contains all clusters). b, Line plots showing scaled expression and gene activities across pseudotime bins of selected examples from multiple K-means clusters. Expression values were smoothed using a generalized additive model. Genes expressed during pluripotency and at the progenitor stage show accurate alignment of the pseudotimes, while neuron specific genes exhibit epigenetic priming (bottom). c, Genome browser snapshots of the neuron specific gene NEUROD2 showing levels of H3K27me3, H3K27ac, H3K4me3 on the gene and RNA expression over pseudotime. d, Quantification of histone modifications and RNA expression on the NEUROD2 locus (c) over pseudotime. e, Jitter plots showing log-normalized fragment counts (for CUT&Tag modalities) and transcript counts (for RNA) along pseudotime. f, Quantification of the shift in pseudotime between the establishment of active histone modifications and RNA expression. g, Detection rate of scATAC and scRNA expression measured from the same cells of a primary developing brain at gestational week 19, the neuron specific genes NEUROD2 and NEUROD6 showing priming of chromatin. h, Gene regulatory network of the neuronal cluster with priming of active epigenetic marks (a - cluster 6) at early (left) and late (right) pseudotime stages. i, Scaled detection rate of upstream regulators (pink - left) and downstream targets (light blue - right) of NEUROD2 over pseudotime. j, Force directed-layout of the telencephalon, diencephalon and mesen-/rhombencephalon trajectory (left). Line plots showing smoothed, averaged pseudotemporal gene expression and activities of k-means clusters with late pseudotime expression in the telencephalon (cluster 1), diencephalon (cluster 1) and mesen-/rhombencephalon (cluster 4) trajectory (right) (see Extended Data Fig. 9a-f for further details). Representative GO-terms of genes within the k-means clusters with late pseudotime expression reveal terms related to neuronal identity (see Supplementary Table 8-10 for details).

We used our single-cell multiome data of the human developing brain (19 gw), where transcription and chromatin accessibility are recorded from the same cell, to test whether neuronal genes show epigenetic priming also in the primary human cortex. Indeed, neuronal genes (NEUROD2, NEUROD6, BCL11B and STMN2) showed a gain in accessibility prior to RNA expression in the developing human cortex (Fig. 3g and Extended Data Figure 8a-e). This supports a role of epigenetic priming during nervous system development, in particular for neuron-specific genes^31,32^ and similar to reports from other developmental systems^6,33–35^. Recently, H3K4me3 has been shown to be required for pausing release of PolII^36^. It is interesting to hypothesize that H3K4me3 marks on these neuronal genes build up gradually before PolII is released into elongation.

To investigate the gene regulatory mechanisms underlying neuronal gene priming, we inferred a gene regulatory network (GRN) based on an integrated scRNA-seq and scATAC-seq dataset of organoid development^14^ informed by histone modification-marked regulatory regions in our scCUT&Tag dataset. We focused on a subnetwork including all genes in the neuronal cluster as well as common transcription regulators that were shared between at least three of the genes in the neuronal cluster. We then summarized epigenetic modifications over pseudotime at transcription factor binding regions. This revealed NEUROD2 as a central hub in this network and showed stepwise activation with loss of repressive marks and gain of active chromatin marks on its regulatory elements throughout pseudotime (Fig. 3h, Supplementary Table 7). We examined the expression of genes acting upstream and downstream of NEUROD2 and found that most upstream regulators (e.g. HEY1, KLF7, SOX2) are expressed after activating marks are installed at the NEUROD2 locus but before NEUROD2 is expressed (Fig. 3i). This suggests that activating chromatin marks establish a transcriptionally permissive landscape prior to NEUROD2 transcriptional regulation. In contrast and as expected, most downstream target genes (like CNTN1, RAS11B) are expressed after activation of NEUROD2 expression (Fig. 3i). To explore epigenetic priming of neuronal genes in other brain region trajectories, we extended the pseudotime analysis to the diencephalon and mesen-/rhombencephalon branch (Fig. 3j). We identified two clusters of late expressed neuronal genes in all branches (e.g. KCNN2, MAP3K13, NEUROD2, SLIT1, STMN2, SLC6A1, NSG1, CPLX2) for which we detected a similar epigenetic priming as observed in the telencephalon (Fig. 3a and j, Extended Data Fig. 9a-f, Supplementary Table 8-10).

Last, we analyzed each gene across all modalities to determine how histone modifications might be differentially involved in brain region diversification versus neuronal differentiation. (Extended Data Fig. 10a-d, Supplementary Table 11). Overall, RNA expression and histone modifications (Extended Data Fig. 10e) showed high concordance. Variance in gene expression and H3K27ac enrichment was mostly explained by regional diversification (Extended Data Fig. 10f and g), whereas H3K27me3 showed similar variance across regional branches and over pseudotime. H3K4me3 showed lower variance across regional branches than over pseudotime (Extended Data Fig. 10g).

Taken together, our analysis provides insight into the dynamic interplay of histone modifications and RNA expression during differentiation from pluripotency to neurons in distinct regions of the developing human brain organoid and shows epigenetic priming of neuronal genes.

### Perturbation experiments show a causal role for histone modifiers in regulating fate acquisition

We next wanted to functionally test the role of histone modifications during early brain organoid development. We chemically inhibited the writer complex of H3K27me3 (EED/PRC2) with A395^37^ during the exit from pluripotency until the formation of the neuroepithelium (day 0-15 of organoid development) to assess the role of EED during this process (Fig. 4a, Extended Data Fig. 11a). Strong developmental defects in full PRC2 KOs have prevented the study of these early brain developmental stages^38,39^. At later developmental stages in mouse cortical development a progenitor-specific depletion of EZH2 altered developmental timing and shifted the balance of progenitor self-renewal and differentiation towards differentiation^40^. We validated the depletion of the H3K27me3 histone marks upon inhibitor treatment by western-blot (Extended Data Fig. 11b). At the neuroepithelial stage (day 15-18) we profiled EED-inhibitor-treated and control (DMSO-treated) organoids using scRNA-seq and bulk CUT&Tag (Fig. 4a and Extended Data Fig. 11c). This data showed a concentration-dependent depletion of H3K27me3 in treated versus control organoids and enrichment of the competitive, activating H3K27ac histone mark at the same sites (Fig. 4b-c, and Extended Data Fig. 11c-d, Supplementary Table 12). Many H3K27me3 peaks also showed co-enrichment for H3K4me3 in the neuroepithelium (Fig. 4b). Next, we used the scRNA-seq data to quantify changes in gene expression linked to H3K27me3 domains upon treatment. We observed a up-regulation of gene expression, in particular for genes in proximity to H3K27me3 depleted peaks (Fig. 4c, Supplementary Table 13). In the highest concentration of EED-inhibitor treatment, nearly all genes neighboring H3K27me3 peaks were significantly up-regulated (Fig. 4c, lower panel). STMN2 and POU5F1 are exemplary genes showing strong H3K27me3 depletion and concomitant gain of H3K27ac and up-regulation of mRNA expression (Fig. 4c-d).

**Fig. 4.**
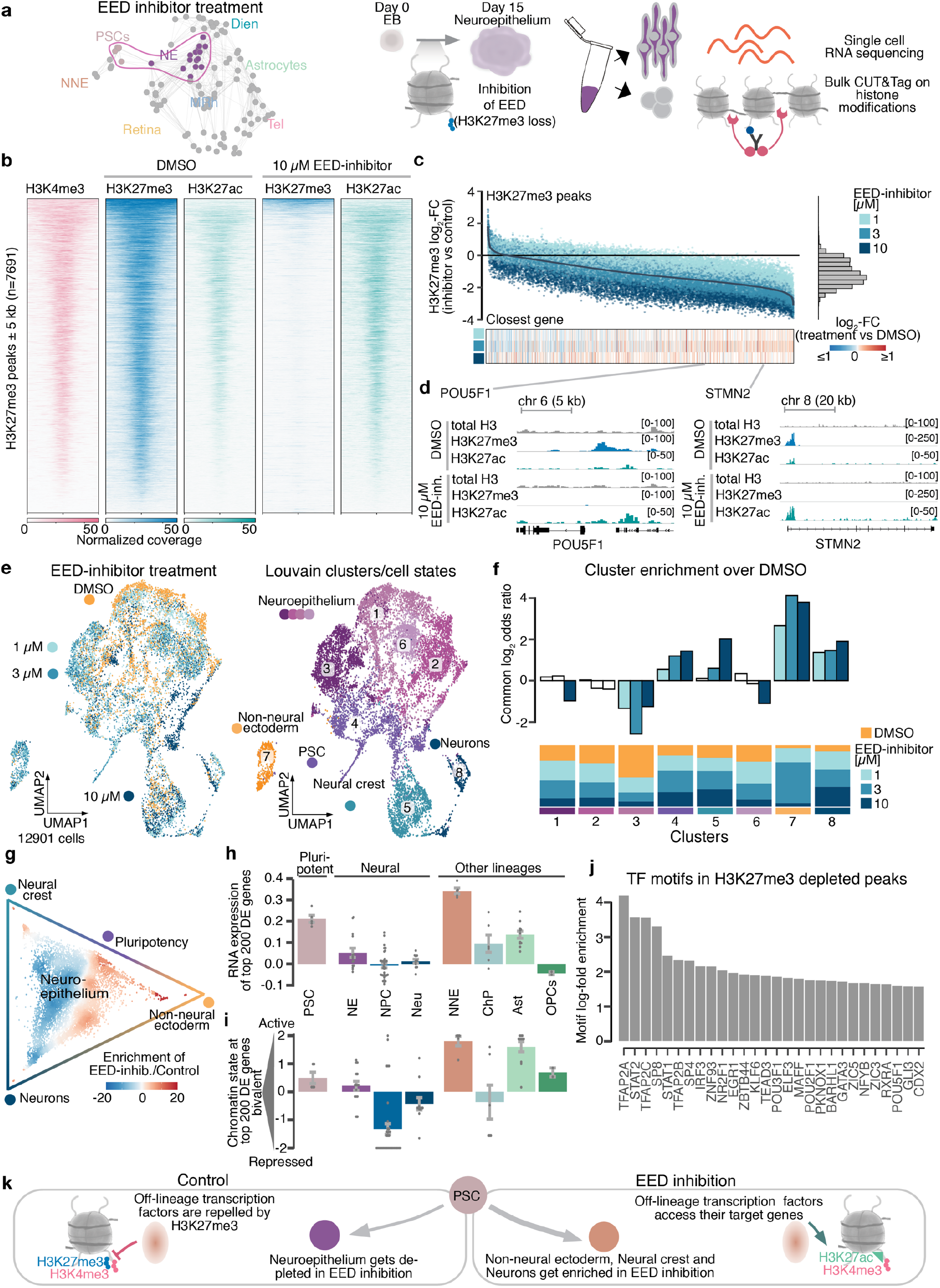
Aberrant cell fate acquisition upon H3K27me3 depletion. a, Schematic of the experiment. Organoids were treated with an EED inhibitor during development from the pluripotent to the neuroepithelium stage (day 0 to day 15). ScRNA-seq and bulk CUT&Tag were performed at the neuroepithelium stage (day 15-18). b, Heatmaps showing H3K4me3, H3K27me3 and H3K27ac bulk CUT&Tag signal intensities on H3K27me3 peaks in DMSO-treated organoids and H3K27me3 as well as H3K27ac signal intensities upon EED-inhibitor treatment (10 µM). Regions are ordered by H3K27me3 intensity. c, Scatter plot showing the log2-fold change of the CUT&Tag signal on H3K27me3-peaks in organoids treated with different EED-inhibitor concentrations versus the DMSO control (top panel) and histogram showing distribution of fold changes (right). Heatmap showing the log2-fold change of the expression of the closest gene from the DE analysis in scRNA-seq data (bottom panel). d, Genomic tracks showing bulk CUT&Tag profiles for H3K27me3 and H3K27ac at genomic regions around STMN2 and POU5F1. e, UMAP embedding of the scRNA-seq data colored by treatment (left) and annotated Louvain clusters (right). f, Bar plot showing cluster enrichment of treated cells versus DMSO control (top) and distribution of treatments in clusters (bottom). Common odds ratio (height) and p-value (alpha) were obtained from a CMH-test stratified by sampling time point. g, Circular plot of differential transition probabilities between the different cell states (e).Inhibitor treated cells show an enrichment at the terminal states of the graph. h, i Barplots quantifying the expression (h) and bivalency (i) of the top 200 DE genes in the cell populations identified from the developmental brain organoid atlas (Fig. 1c). j, Transcription-factor motif enrichment in H3K27me3 depleted peaks close (10 kb) to differentially expressed genes in cluster 7 (see Extended Data Fig. 12e-f for details). k, Schematic showing how H3K27me3 mediated repression of transcription factor motifs could affect lineage decisions when cells exit from pluripotency and lead to preferential stabilization of the non-neural ectoderm fate.

To assess how such strong changes in transcriptome and epigenome impact cell identities, we integrated the scRNA-seq data between conditions, performed clustering and annotated the resulting populations (Fig. 4e). As expected from this time point, we observed a large neuroepithelial progenitor population (cluster 1, 2, 3, 6), however we observed an enrichment of inhibitor-treated cells in populations of pluripotent cells (cluster 4) non-neural ectoderm (cluster 7), neural crest (cluster 5) and neurons (cluster 8) (Fig. 4e-f and Extended Data Fig. 11e-g). We used CellRank^22,41^ to calculate transition probabilities into clusters resembling terminal states (non-neural ectoderm, neural crest and neurons) and confirmed that cells transition from the neuroepithelium and pluripotency state to non-neural ectoderm, neural crest and neurons - off-lineage cell states enriched upon inhibitor treatment (Fig. 4g and Extended Data Fig. 11h). We performed differential expression analysis in the neuroepithelium cluster and found that many treatment-upregulated genes were also marker genes for the ectopic clusters (Extended Data Fig. 12a, ANXA2, S100A10, non-neural ectoderm; STMN2, NEFM, neurons, Supplementary Table 14-15). These genes exhibit an inhibitor-dependent loss of H3K27me3, gain in H3K27ac and upregulation of mRNA (Extended Data Fig. 12b), indicating that loss of H3K27me3 shifted cell identity towards ectopic cellular states. Indeed, the top upregulated genes upon H3K27me3 inhibition were expressed outside of the neuroepithelium in our developmental atlas, predominantly in pluripotency, non-neural ectoderm and astrocytes (Fig. 4h). We found that genes most strongly upregulated upon loss of H3K27me3 resided in H3K27me3/H3K4me3 bivalent domains within the neuroepithelium and PSCs, thereby primed for activation upon H3K27me3 loss (Fig. 4b, i). This suggests that H3K27me3 is required to stabilize cell fate decisions when cells progress from pluripotency to neuroepithelium. Loss of H3K27me3 at the pluripotency and neuroepithelium stage mostly affects genes that are in a permissive chromatin environment (H3K27me3/H3K4me3). These genes become activated thereby inducing the cell to transition to a cell state that would normally be marked by these genes, such as non-neural ectoderm or pluripotency (Fig. 4h and i, Extended Data Fig. 12c-d). This suggests that removal of H3K27me3 causes primarily upregulation of genes within transcriptionally permissive regions.

We investigated which transcription factors might be involved in destabilizing cell fate decisions at the exit from pluripotency by analyzing binding motifs within H3K27me3 depleted peaks in close proximity to upregulated genes in the ectopic clusters (Fig. 4j). This revealed a motif enrichment for the TFAP2-family members that regulate specification of neural crest and non-neural ectoderm^42,43^, STAT-family members that are involved in cell proliferation, and multiple transcription factors that regulate neuronal developmental processes (e.g. EGR1, KLF6 and SP8). GO term analysis on these transcription factors with motif enrichments revealed their involvement in neuron apoptotic processes, differentiation and organ development (Extended Data Fig. 12e). Most of these transcription factors are expressed during the neuroepithelium stage (Extended Data Fig. 12f). We hypothesize that these transcription factors have enhanced access to their target genes upon depletion of H3K27me3 and can induce a cascade of dysregulation that ultimately alters cell fate choice and leads to enrichment of ectopic fates such as neural crest and non-neural ectoderm or accelerated fates that would occur at later developmental stages like neurons (Fig. 4k). Our observation offers a mechanistic explanation to the alterations in developmental timing observed in other systems^10,40,44^. Interestingly, around 60% of cells still established neuroectodermal and neuroepithelial identities, even in the nearly complete absence of H3K27me3. This further supports that the neuroectoderm comprises a default fate that cells assume in the absence of further inductive signals^45,46^.

Overall, we identified the molecular framework of how H3K27me3 mediated gene repression ensures lineage fidelity during early central nervous system development. We delineate at single cell resolution its effect on immediate branch points within early developmental stages.

## Discussion

During early human brain development, gene expression must be tightly coordinated to enable controlled differentiation into a multitude of cell types. So far, it has remained challenging to interrogate the epigenetic mechanisms that shape these dynamic processes on a global scale and with single-cell resolution. Here we have addressed this challenge by measuring histone modifications at single-cell resolution in brain organoids, which model important aspects of early human brain regionalization and cell type establishment *in vitro*. Over a developmental organoid time course, we integrated scCUT&Tag profiles marking repressed and active chromatin states with RNA-expression data. Broadly, we found that chromatin modification profiles were highly specific for a given cell population. We identified region-specific regulatory elements in close proximity to important cell fate determinants. Such determinants were also often marked by epigenetic switches, which shift from repressive to active chromatin states at fate bifurcation events to prime gene expression. These observations indicated that chromatin modifiers play crucial roles in regional diversification events and stabilize cell fate commitment. This was further supported by perturbation of EED, which caused a global depletion of H3K27me3, and up-regulation of an otherwise repressed but poised set of genes and a strong tendency of cells to adopt off-target cell states. Altogether, these results suggest that dynamic histone modifications are required to ensure robust cell fate acquisition and brain regionalization. Moreover, our developmental atlas of histone marks will serve as a reference to better understand the epigenomic landscape of human brain development and will be instructive for future studies on cell fate commitment and reprogramming.

## Materials and Methods

### Cell and organoid culture

Cell lines used in the study were derived from different sources (Method Table 1, below).

**Method Table 1:** Cell lines used in the study.

| Cell line | Sex | Origin |
| --- | --- | --- |
| 409b2-iCRISPR | female | originally from the RIKEN BRC cell bank modified S. Pääbo lab<sup>47</sup> |
| HOIK1 | female | HipSci resource<sup>48</sup> |
| WIBJ2 | female | HipSci resource<sup>48</sup> |
| WTC (GM25256) | male | Coriell Institute<sup>49</sup> |
| B7 | female | B. Roska lab<sup>50</sup> |

For culturing cells were grown on matrigel (Corning, #354277) coated 6-well dishes in mTeSR Plus (StemCell Technologies, #100-0276) supplemented with penicillin/streptomycin (P/S, 1:200, Gibco, #15140122). To propagate the cells, they were dissociated with TryplE (Gibco, #398 12605010) or EDTA in DPBS (final concentration 0.5mM) (Gibco, #12605010) and kept on Rho-associated protein kinase (ROCK) inhibitor Y-27632 (final concentration 5 μM, StemCell Technologies, #72302) for one day. Cells were stored in liquid nitrogen in mFreSR (StemCell Technologies, #05855) and tested for mycoplasma (Venor GeM Classic, Minerva Biolabs) after each thawing cycle.

To generate cerebral organoids, cells were grown to a confluency of around 50% and dissociated with TryplE. 2000-3000 cells were aggregated in 96-well ultra low attachment plates (Corning, #CLS7007) to form embryoid bodies (EBs). We followed an unguided protocol to obtain brain organoids^51^, with a few modifications. EBs were aggregated and cultured in mTeSR Plus and neural induction medium was added when the EBs had reached around 400-500 µm (usually day 5). Retinoic acid containing neural differentiation medium was only added from day 40 onward^32^. Cerebral organoids were grown shaking in 6 cm dishes for up to 8 months. To generate retinal organoids, we applied a protocol that allows the simultaneous aggregation of hundreds of embryoid bodies in agarose molds^50^. After neural induction, these are transferred (1 week) to matrigel coated 6-well plates and allowed to attach. The neural induction medium used in both protocols is highly similar allowing to comparability of the neuroepithelium stage (both media contain DMEM / F12 (GIBCO, #31331‒028), 1xN2 Supplement (GIBCO, #17502‒048), 1% NEAA Solution (Sigma, #M7145) and 2 mg/ml heparin (Sigma, #H3149‒50KU) (1µl/ml for brain organoids), in case of brain organoids 1% Glutamax was added). After 2 weeks, neural differentiation medium is added and retinal structures are scraped off after 4 weeks.

### Drug treatment

We used a pharmacological approach to inhibit chromatin modifiers in cerebral organoids. Specifically we targeted the H3K27me3-reader EED^37^ (A395, MedChemExpress #HY-101512) that prevents allosteric activation of the PRC2 complex. For A395 concentrations between 1-10 µM were tested and H3K27me3 was depleted in bulk experiments at 1 µM as shown by western blot. EED was inhibited from day 0-15 (experiment 1) and day 0-18 (experiment 2), by adding the inhibitor when seeding the EBs. This time window coincides with the formation of the neural epithelium. Following, the medium was changed every other day adding a fresh dose of inhibitors. To rule out cell line specific effects we treated two different cell lines (HOIK1 and WIBJ2). We used single nucleotide polymorphisms (SNPs) to assign the cells bioinformatically to each cell line.

### Preparation of single cell suspensions

Brain organoids were generated in batches. In each batch five different stem cell lines (409b2-iCRISPR, B7, WTC, HOIK1 and WIBJ2) were used and dissociated together. The cell lines were later demultiplexed using single nucleotide polymorphisms (SNPs). For retinal organoids only B7 was used. We sampled all important developmental transitions from the pluripotent EB (day 5) and neuroepithelium (day 15) to neuronal differentiation in brain (day 35, 60, 120, 240) and in retina organoids (day 45 and 85). Organoids were cut into small pieces using a scalpel and thoroughly washed with HBSS buffer without Ca2+/Mg2+ (StemCell Technologies, #37250) to remove debris.

To obtain single cell suspensions a papain-based neural dissociation kit (Miltenyi Biotec, 130-092-628) was used. In brief, 1900 µl of pre-warmed buffer X with 50 µl of Enzyme P were added to the organoids and incubated for 15 min at 37 °C. Subsequently, a mix of 20 µl buffer Y and 10 µl of DNase were added to each sample before tituration (10 times with a p1000 wide bore tip, 10 times with a p1000 wide bore tip). The samples were then incubated twice for another 10 min and titurated with a p1000 and p200 in between.

At the end, the reaction was stopped with HBSS buffer without Ca2+/Mg2+ and the cells were passed through a 30 µm strainer. After an additional wash the cells were then stored in CryoStor CS10 (StemCell Technologies, Catalog # 07930) or processed right away for further experiments. For all scCUT&Tag experiments, cells were kept from the same cell suspension to perform scRNA sequencing. Cell suspensions usually exhibited viabilities between 80-95%.

### Preparation of single cell suspensions from human fetal brain

Human fetal brain tissue (19 gestational weeks) was obtained following elective pregnancy termination and informed written maternal consents from Advanced Bioscience Resources, CA, USA. The estimated age of the fetus was calculated using clinical information like the last menstrual period and anatomical data obtained through ultrasound measurements. Dissected fetal brains were kept in DMEM+antibiotics on wet ice until preparation of the single cell suspension for up to 48 hrs. Single cell suspensions were prepared following the same protocol as for organoids. At this developmental time point, the cortex has already expanded and makes up for the majority of the cells in the population. We tried to enrich cortical material by separating larger pieces from the material. The resulting single cell suspensions were cryopreserved until further use. Nuclei suspensions for 10x multiome experiments were prepared following the single cell CUT&Tag protocol and the nuclei were processed following the manufacturers’ instructions.

### Cloning and purification of Tn5

Plasmids #123461 (pA/G-MNase) and #124601 (3XFlag-pA-Tn5-Fl) were ordered from Addgene. ProteinA and ProteinG were amplified using the primer pairs (FZ461_ProtA_rev, FZ462_HindIII_ProtA_fw and FZ459_EcoRI_ProtG_rev, FZ460_ProtG_fw) and fused by PCR (all primer sequences are in Method Table 2, below). The ProteinA in the original vector (3XFlag-pA-Tn5-Fl, addgene #124601^52^) was then replaced with the fusion product through EcoRI and HindIII restriction digest.

The final plasmid was transformed into chemically competent Rosetta cells to express the protein. The bacteria were grown to OD_600_∼0.4-0.6, expression was induced with 0.25 mM IPTG and the protein was expressed at 18 °C overnight. Cells were harvested and stored at - 80 °C until further processing. The purification was performed on Chitin resin (New England Biolabs, #S6651S) as described^53^, with small modifications. The cells were lysed using the Diagenode Bioraptor Plus at the high setting for 15 cycles 30 sec on/30 sec off.

After dialysis and concentration using Amicon Ultra-4 Centrifugal filters (Millipore, #UFC803024) the protein was diluted to 50% glycerol final and loaded with adapters before use (Method Table 2, below).

**Method Table 2:**
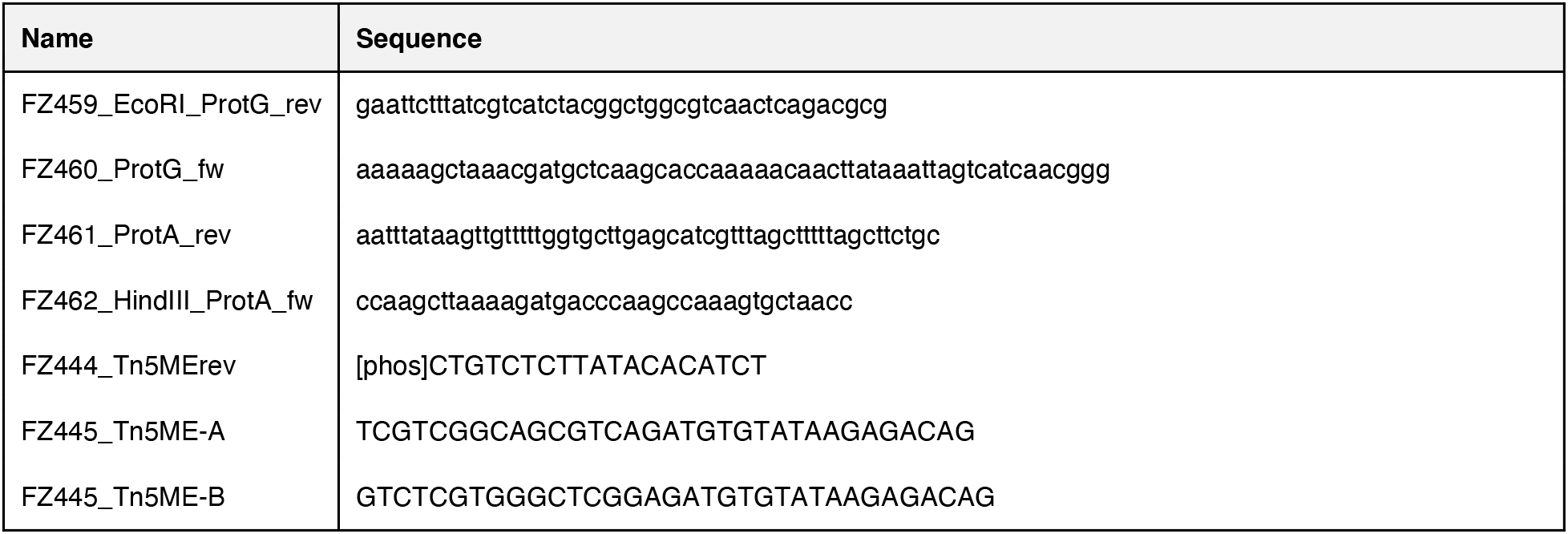
Primers used in the study.

### Single cell CUT&Tag

Starting from 1.5-3 Mio cells, nuclei were isolated following the 10x Genomics CG000365 Demonstrated Protocol. The 0.1x buffer was used for all experiments, adjusting the final concentration of Digitonin to 0.01% (Thermofisher, BN2006). In general, we followed the bulk CUT&Tag protocol with a few adjustments^52^.

After lysis the nuclei were directly transferred into scCUT&Tag wash buffer (20 mM HEPES [pH 7.5] (Jena Bioscience, #CSS-511), 150 mM NaCl (Sigma Aldrich, #S6546), 0.5 mM Spermidine (Sigma Aldrich, #S0266), 1% BSA (Miltenyi, # 130-091-376), 1 mM DTT (Sigma Aldrich, #10197777001), 5 mM sodium butyrate (Sigma Aldrich, #303410), Roche Protease Inhibitor (Sigma Aldrich, #11873580001)) and washed once for 3 min at 300 xg. The buffer was supplemented with 2 mM of EDTA before the antibodies were added (see Method Table 3 for further details). The samples were incubated on a rocking platform at 4 °C overnight. The next morning the cells were washed once and the secondary antibody was added to the suspension for 1 hr at 20 °C on a rocking platform or Eppendorf Thermomixer. The cells were then washed twice and transferred into scCUT&Tag med buffer (20 mM HEPES [pH 7.5] (Jena Bioscience, #CSS-511), 300 mM NaCl (Sigma Aldrich, #S6546), 0.5 mM Spermidine (Sigma Aldrich, #S0266), 1% BSA (Miltenyi, # 130-091-376), 1 mM DTT (Sigma Aldrich, #10197777001), 5 mM sodium butyrate (Sigma Aldrich, #303410), Roche Protease Inhibitor (Sigma Aldrich, #11873580001)) including 2 µg of homemade Tn5. The cells were incubated for 1 hr at 20 °C and then washed again twice with scCUT&Tag med buffer. To induce the cutting of the Tn5, 10 mM final MgCl_2_ (Sigma Aldrich, #M1028) was added and the sample was incubated for 1 hr at 37 °C. After the incubation, the reaction was stopped with 15 mM EDTA final and the sample was filled up to 600 µl with diluted nuclei buffer of the 10x Genomics scATAC kit v1.1 supplemented with 2% BSA. The nuclei were filtered through a 40 µm Flowmi filter (Sigma Aldrich, #BAH136800040) and washed with diluted nuclei buffer supplemented with 2% BSA. The final nuclei suspension was quality controlled and counted with a Trypan Blue assay on the automated cell counter Countess (ThermoFisher). Finally 15-20k nuclei were loaded per experiment. In cases where fewer nuclei were recovered all nuclei were loaded. To run the Chromium Chip, 5 µl of cell suspension were mixed with 3 µl of PBS and 7 µl of ATAC buffer from the kit.

Libraries were prepared following the manufacturer’s instructions, except that 2 PCR cycles were added to the barcoding PCR and after 10 cycles of indexing PCR 5 µl of the library were used to determine the final number of cycles in a Roche LightCycler^54^. Usually sCUT&Tag libraries required 12-16 PCR cycles during the indexing. After SPRI select clean-up (Beckman Coulter, #B23318) the libraries were quality controlled and sequenced following the 10x Genomics scATAC v1.1 sequencing recommendations. Usually 50-100 Mio reads per library were enough to cover the complexity.

**Method Table 3:**
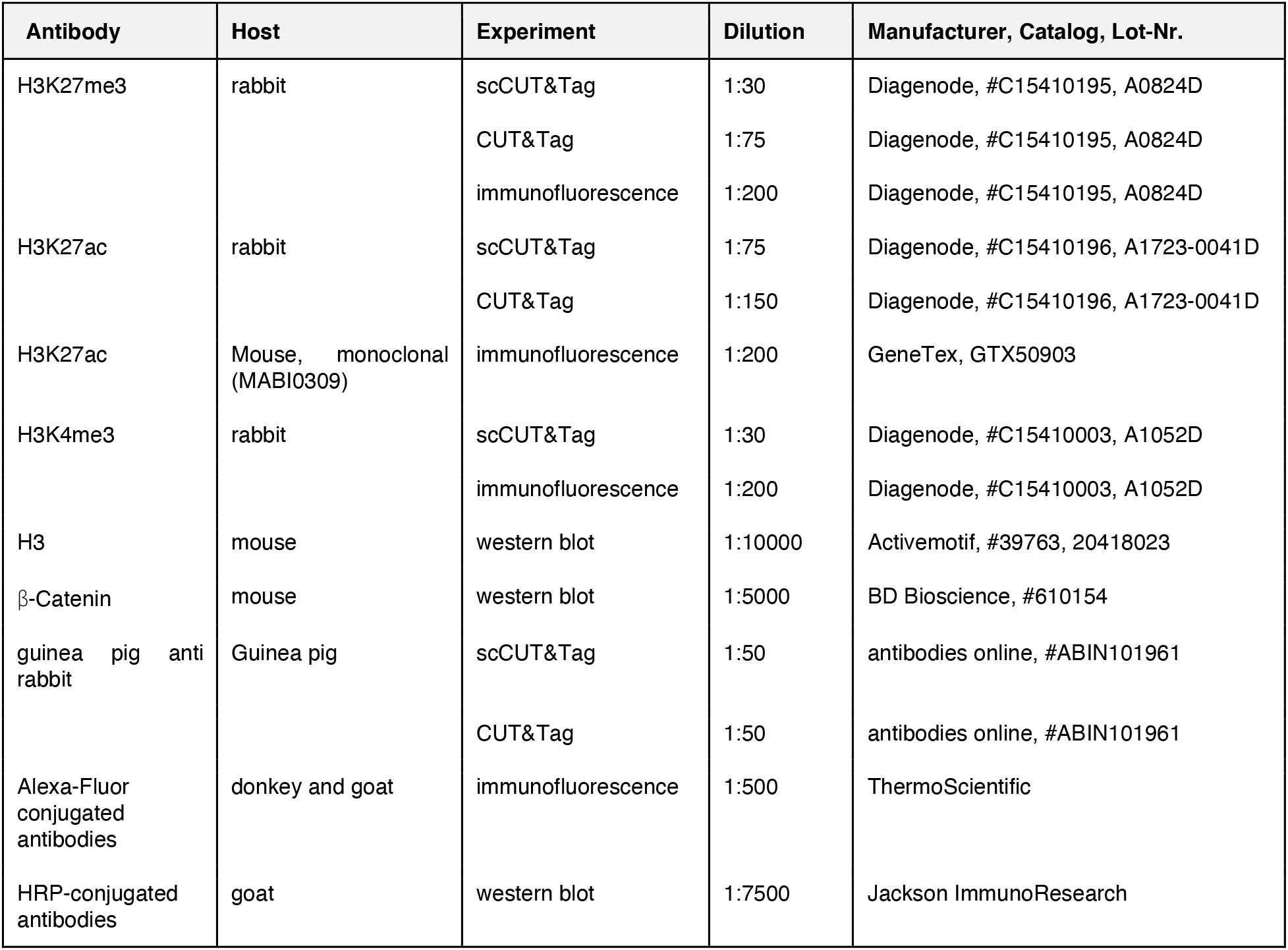
Antibodies used in the study.

### Bulk CUT&Tag

Starting with 0.1-1 Mio cells after dissociation, cells were transferred into CUT&Tag wash buffer (20 mM HEPES [pH 7.5] (Jena Bioscience, #CSS-511), 150 mM NaCl (Sigma Aldrich, #S6546), 0.5 mM Spermidine (Sigma Aldrich, #S0266), 5 mM sodium butyrate (Sigma Aldrich, #303410), Roche Protease Inhibitor (Sigma Aldrich, #11873580001)). Following 15 µl of BioMag ConcanavalinA beads (Polysciences, #86057-3) in binding buffer (20 mM HEPES (pH 7.5), 10 mM KCl, 1 mM CaCl_2_, 1 mM MnCl_2_) were added to the sample and incubated on the wheel for 15 min at RT. Subsequently the cells were collected on a magnet and lysed through the addition of CUT&Tag wash buffer supplemented with 0.01% Digitonin. Lysis was monitored under a microscope with Trypan Blue staining. After lysis was complete the nuclei were washed again with CUT&Tag wash buffer. If possible all samples were split and H3 or another chromatin mark CUT&Tag was performed on the same starting material, to be used as normalizer. The antibody was added together with 2 mM EDTA final and the sample was incubated on a rocking platform at 4 °C overnight.

The samples were washed once with CUT&Tag wash buffer and the secondary antibody was added to the reaction and incubated for 1 hr at 20 °C on a rocking platform. After two additional washes the Tn5 was added (1:100) in CUT&Tag med buffer (20 mM HEPES [pH 7.5] (Jena Bioscience, #CSS-511), 300 mM NaCl (Sigma Aldrich, #S6546), 0.5 mM Spermidine (Sigma Aldrich, #S0266), 5 mM sodium butyrate (Sigma Aldrich, #303410), Roche Protease Inhibitor (Sigma Aldrich, #11873580001)). Tn5 was allowed to bind for 1 hr at 20 °C on a rocking platform. After two additional washes the cutting was induced through addition of 10 mM MgCl_2_ in CUT&Tag med buffer. After 1 hr at 37 °C the reaction was stopped by adding a final of 20 mM EDTA, 0.5% SDS and 10 mg Proteinase K. The reaction was then incubated at 55 °C for 30 min and finally inactivated at 70 °C for 20 min.

The DNA fragments were purified using the ChIP DNA Clean & Concentrator kit (Zymo Research, #D5205) for the elution from the columns 2 pg of Tn5-digested and purified lambda DNA (New England Biolabs, # N3011S) were added to be used as spike-in normalizer for later analysis, when needed. Purified fragments were indexed for 15 cycles (1 x 5 min at 58 °C, 1 x 5 min at 72 °C, 1 x 30 s at 98°C, 14x 10 s at 98 °C, 30 s at 63 °C, 1 x 1 min at 72 °C, ∞ at 4 °C) using NEBNext HighFidelty 2x PCR Master Mix (New England Biolabs, M0541S) and Illumina i5 and i7 indices^54^. The libraries were then purified using AmPure beads (Beckman Coulter, #A63881), measured and quality controlled with Qubit DNA HS Assay (ThermoScientific, #Q32854) and on the Tapestation (Agilent, #5067-4626) and then sequenced (PE, 2×50 bp).

### Hashing and single cell RNA-Seq

Cells were either processed right after dissociation or recovered after cryostorage. To recover the cells after cryostorage, the cryovials were incubated in a water bath at 37 °C until only a small ice-piece was left inside the tube. The cells were then transferred into prewarmed DMEM/F-12 (Gibco, #31330038) supplemented with 10% FBS final (Merck, #ES-009-B). After washing twice with DPBS (Gibco, # 14190144) supplemented with 0.5% BSA the cells were filtered with a 40 µm Flowmi filter and counted using the Trypan Blue assay on the automated cell counter Countess (ThermoFisher). Usually cell viability was around 80-95%. At this point cells were further diluted to be processed in the 10x Genomics single cell RNA expression v3.1 assay following strictly the manufacturer’s guidelines.

To generate single cell RNA expression libraries of the drug treatment we turned to cell hashing^55^ for better comparability of the samples. For hashing 100-300 k cells were resuspended in 50-100 µl of DPBS+0.5% BSA. Following 5 µl of the Human TruStain FcX (Fc Receptor Blocking Solution, BioLegend, #422302) were added to the sample and incubated for 10 min on ice. Next, the 2 µl TotalSeq Cell hashing antibodies (BioLegend) were added to the cells and incubated for 30 min on ice with gentle agitation every 10 min. After incubation cells were washed twice with DBPS+0.5% BSA. Depending on the starting amount the cells were resuspended in 20-40 µl DPBS+0.5% BSA and counted. Following the cell suspensions were mixed and processed in the 10x Genomics single cell RNA expression v3.1 assay following strictly the manufacturer’s guidelines with slight adjustments. A maximum of 20 k cells was targeted per experiment and HTO additive primer was added to the cDNA synthesis following the TotalSeq technical protocol (https://www.biolegend.com/en-us/protocols/totalseq-a-antibodies-and-cell-hashing-with-10x-single-cell-3-reagent-kit-v3-3-1-protocol). Togenerate gene expression and hashing libraries we followed the protocol CG000206 Chromium Next GEM SingleCell v3.1 Cell Surface Protein and sequenced according to the manufacturer’s guidelines.

### Western Blot

Depending on the time point 1-3 organoids were directly collected into 50 µl Laemmli sample buffer and homogenized with an electric grinder (Fisherbrand 12-141-368). DNA was sheared by sonication in the Diagenode Bioraptor Plus (high setting for 15 cycles 30 sec on/30 sec off). Samples were subsequently run on SDS-PAGE and transferred to PVDF membrane using Wet-Blot. 2-10 µl of extract were loaded per lane. The ECL signal was recorded using the iBright system (Invitrogen). The signal was compared to antibody stainings of loading controls (H3 and β-Catenin) and the membranes were quality controlled by Ponceau (Sigma Aldrich, #P7170-1L) staining. See Method Table 3, above, for details on the antibodies.

### Immunostaining

Organoids were fixed overnight in 4% PFA. The next day, the organoids were washed three times for 5 min with DPBS and then transferred into 30% sucrose in DPBS until they sank to the bottom of the tube. Following the organoids were transferred into cryomolds (Sakura, #4565) and embedded in Tissue-Tek O.C.T. (Sakura, #16-004004) on dry ice. The organoids were then sliced at the Cryostar NX70 (Thermo Scientific) into 20 µm thick slices at −17 °C. The slices were transferred to glass slides and washed with PBS. After a quick wash, antigen retrieval was performed for 20 min at 70 °C in 1x preheated HistoVT one (Nacalai, #06380). Slides were washed 3 times for 5 min with PBS+0.2% Tween and then transferred to blocking and permeabilization (PBS, 0.1% Triton, 5% Serum, 0.2% Tween, 0.5% BSA) for 1 hr. The antibodies were added in blocking solution overnight at 4 °C (see Method Table 3, above, for further details). The next day the slides were washed again 3 times for 5 min with PBS+0.2% Tween and the secondary antibody was added in PBS supplemented with 2% BSA and 0.2% Tween for 2 hrs at room temperature. Last, the slides were washed again 3 times for 5 min with PBS+0.2% Tween adding DAPI to the last wash. The slides were then mounted in Prolong Glass Antifade and imaged at the Nikon Ti2 Spinning Disk or at the confocal Zeiss LSM980.

### Data processing for scRNA-Seq

To compute transcript count matrices sequencing reads were aligned to the human genome and transcriptome (hg38, provided by 10x Genomics) by running Cell Ranger (version 5.0.0) with default parameters. Count matrices were then preprocessed using the Seurat R package (version 3.2)(*41*).

Cells were filtered by unique molecular identifier (UMI), number of detected genes, and fraction of mitochondrial genes as follows:

UMIs > 2000

UMIs < 1.5 * 10^5^

detected genes > 1000

fraction of mitochondrial reads < 0.2

Transcript counts were normalized by the total number of counts for that cell, multiplied by a scaling factor of 10,000 and subsequently natural-log transformed (NormalizeData()).

### Preprocessing and clustering of scCUT&Tag data

We aligned the sequencing reads to the human genome and transcriptome (hg38, provided by 10x Genomics) using Cell Ranger ATAC (version 1.2.0) with default parameters to obtain fragment files and peak calls. The fragment files and the peak count matrices were further preprocessed using Seurat (version 3.2)^56^ and Signac (version 1.1)^57^. We removed cells with less than 200 (H3K27ac, H3K4me3) or 100 (H3K27me3) fragment counts from the analysis. For quality control we checked the following metrics using Signac: The transcription start site (TSS) enrichment score (TSSEnrichment()), in particular for activating and promoter marks, the, nucleosome signal (NucleosomeSignal()), the percentage of reads in peaks, and the ratio of reads in genomic blacklist regions.

We then created a unified set of peaks from the union of peaks from all samples by merging overlapping and adjacent peaks. The unified set of peaks was re-quantified for each sample using the fragment file (FeatureMatrix()). Peak counts were normalized by term frequency-inverse document frequency (tf-idf) normalization using the Signac functions RunTFIDF(). Latent semantic indexing (LSI) was performed by running SVD (RunSVD()) on the tf-idf-normalized matrix. To visualize the data in 2D, Uniform Manifold Approximation and Projection (UMAP)^12^ was performed on LSI components 2-30. We then called high-resolution clusters using Louvain clustering in each group separately with the following resolutions to obtain similar cluster sizes:

EB: 2

mid: 5

late: 10

8 months: 5

retina: 5

Demultiplexing

We used demuxlet^58^ to demultiplex cells pooled from different stem cell lines. For B7 and 409B2-iCRISPR single nucleotide polymorphisms were called using bcftools based on DNA-seq^47,59^ data or downloaded from the HipSci website (HOIK1, WIBJ2) and the Allen Cell Atlas (WTC). All files were merged using bcftools and sites with the same genotypes in all samples were filtered out. Demuxlet was run with default settings. For RNA, cells with ambiguous or doublet assignments were removed from the data. Otherwise, the best singlet assignment was considered the lines’ genotype.

## Integration and annotation of scRNA-seq data

First, we grouped the dataset into 5 groups depending on the sample of origin:

EB: 4 days old brain organoids in EB stage

mid: 15 days old brain organoids in neuroepithelium stage late: 1-4 month old brain organoids

8 months: 8 month old brain organoids

retina: 6 week & 12 week old retina organoids

Initial integration was based on mid and late groups. We computed the 2000 most variable features using the Seurat function FindVariableFeatures() and computed cell cycle scores using the Seurat function CellCycleScoring(). Subsequently the data was z-scaled, cell cycle scores were regressed out (ScaleData()) and Principal Component Analysis (PCA) was performed using the Seurat function RunPCA() based on variable features. We used the first 10 principal components (PCs) to integrate the different time points in the dataset using the Cluster Similarity Spectrum method (CSS)^60^. The missing samples EB, retina and 8 months were then projected into CSS space using the css_project() function. To obtain a two-dimensional representation of the data we performed Uniform Manifold Approximation and Projection (UMAP)^12^ using RunUMAP() with spread=0.8, min.dist=0.2 and otherwise default parameters.

To annotate the data, we first called high-resolution clusters using Louvain clustering in each group separately with the following resolutions to obtain similar cluster sizes:

EB: 1

mid: 2

late: 5

8 months: 2

retina: 2

Clusters were annotated with cell types and regional identities using VoxHunt^61^, comparison to reference datasets^13,14^ and marker expression.

### Hashing

Count matrices were generated with CITE-seq-Count (version 1.4.5 running under Python version 3.6.13) and intersected with transcript matrices from Cell Ranger. The hashtag counts were normalized with centered log-ratio (CLR) transformation. Doublets were filtered out using hashtag information.

### Calculation of gene activity scores

To enable comparison of gene expression with chromatin modifications in the same feature space, we computed gene activity scores for each gene and chromatin modality. For this we used the Signac function GeneActivity() with default parameters. Fragment counts representing gene activities were subsequently log-normalized with a scaling factor of 10,000.

### Matching of scRNA and scCUT&Tag

To integrate cell populations between RNA and chromatin modalities, we matched high-resolution clusters based on correlation of gene expression with gene activity scores. For this, we performed minimum-cost maximum-flow (MCMF) bipartite matching between the modalities as described in (https://github.com/ratschlab/scim)^21^. The function get_cost_knn_graph() was used with knn_k=10, null_cost_percentile=99 and capacity_method=’uniform’. As a distance metric (knn_metric), we used the correlation distance provided by scipy^62^ for activating marks H3K27ac and H3K4me3 and the negative correlation distance for the repressive mark H3K27me3. Unmatched clusters from either modality were matched based on maximum (or minimum) correlation.

### Graph representation of regional diversification

To visualize differentiation trajectories into regionalized neuronal populations, we constructed a graph representation based on terminal fate probabilities. For this, we first obtained count matrices for the spliced and unspliced transcriptome using kallisto (version 0.46.0)^63^ by running the command line tool loompy fromfastq from the python package loompy (version 3.0.6)(https://linnarssonlab.org/loompy/). We subset the dataset to cell populations on the neuronal trajectory from pluripotency (PSCs, neuroectoderm, NPCs and neurons) and computed RNA velocity using scVelo (version 0.2.4)^22^ and scanpy (version 1.8.2)^64^. First, 2000 highly variable features were selected using the function scanpy.pp.highly_variable_genes(). Subsequently, moments were computed in CSS space using the function scvelo.pp.moments() with n_neighbors=20. RNA velocity was calculated using the function scvelo.tl.velocity() with mode=’stochastic’ and a velocity graph was constructed using scvelo.tl.velocity_graph() with default parameters. To order cells in the developmental trajectory, a root cell was chosen randomly from cells of the first time point (EB) and velocity pseudotime was computed with scvelo.tl.velocity_pseudotime(). The obtained velocity pseudotime was further rank-transformed and divided by the total number of cells in the dataset. Based on the velocity pseudotime, we computed fate probabilities into the following manually annotated terminal cell states: Dorsal telencephalon neurons, diencephalon neurons, midbrain neurons, hindbrain neurons and retinal ganglion cells. For this, we used CellRank (version 1.3.0)^41^. A transition matrix was constructed with a palantir kernel (PalantirKernel()) based on velocity pseudotime. Absorption probabilities for each of the predefined terminal states were computed using the GPCCA estimator. Based on the computed fate probabilities we next constructed a graph representation. We used PAGA to compute the connectivities between clusters (scvelo.tl.paga()) and summarized transition scores for each of the clusters. To find branch points at which the transition probabilities into different fates diverge, we constructed a nearest-neighbor graph between the high-resolution clusters based on their transition scores (k=10). We further pruned the graph to only retain edges going forward in pseudotime, i.e. from a node with a lower velocity pseudotime to a node with a higher velocity pseudotime. Additionally, we removed edges connecting different regional trajectories. The resulting graph is directed with respect to pseudotemporal progression and represents a coarse-grained abstraction of the fate trajectory, connecting groups of cells with both similar transition probabilities to the different trajectories and high connectivities on the transcriptomic manifold.

### Differential peak analysis between regional identities

To find peaks with differential enrichment in regional trajectories, we performed differential peak analysis for each chromatin modality. We fit a generalized linear model with binomial noise and logit link for each peak i on binarized peak counts Y with the total number of fragments per cell and the region label as the independent variables:

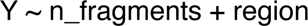

In addition, we fit a null model, where the region label was omitted:

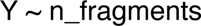

We then used a likelihood ratio test to compare the goodness of fit of the two models using the lmtest R package (version 0.9) (https://cran.r-project.org/web/packages/lmtest/index.html). Multiple testing correction was performed using the Benjamini-Hochberg method.

### Analysis of bivalent and switching peaks

To find genomic regions that were marked by both H3K27me3 and H3K4me3 (bivalency), we extended all peaks of both modalities by 2 kb in both directions before intersecting them. For all intersecting peaks, we aimed to find instances where bivalency is resolved during regional diversification. For this, selected matching peaks that showed differential enrichment in any region with opposite effect sizes. Out of these, we further selected matching peaks where both modalities were detected in >5% of cells in any high-resolution cluster in the neuroepithelial stage (bivalent in neuroepithelium). We performed an analogous analysis for H3K27me3 and H3K27ac to find regions where switching occurs upon regional diversification. Here, we selected matching peaks from both modalities for which only one mark was detected in the neuroepithelial stage (>5% detection in any high-resolution cluster).

### Reconstruction of the telencephalic neuron differentiation trajectory from pluripotency

To reconstruct the differentiation trajectory leading up to telencephalic neurons in higher resolution, we first extracted all cells annotated as EB, neuroepithelium, telencephalic progenitors and neurons. We next sought to compute a pseudotime describing the progression along this trajectory for all modalities separately. For all chromatin modalities, we used LSI components 2-10 to compute diffusion maps with the R package destiny^30^. Ranks along the first diffusion component were used as a pseudotemoral ordering. For RNA, we used the function scvelo.tl.velocity_pseudotime() from scVelo^22^ to compute a pseudotime based on RNA velocity. To obtain an even distribution of time points for all modalities, we next subsampled the trajectory for each time point group to the lowest cell number in any modality but a minimum of 100. The subsampled trajectory was then stratified into 50 bins of equal cell number or 20 bins in case of neurogenesis trajectories.

### Clustering of pseudotemporal expression patterns

To discover groups of genes with similar pseudotemporal expression patterns, we clustered smoothed expression patterns based on a dynamic time warping distance. For this, genes for clustering were selected by intersecting 6000 highly variable genes in RNA with genes detected in >2% of cells in any pseudotime bin for all chromatin modalities. For these genes, the average log-normalized expression and gene activity was computed for each modality and pseudotime bin. We smoothed the mean expression for each gene’s values using a generalized additive model with a cubic spline (bs=’cs’), which was fit using the R package mgcv^65^. Smoothed expression over pseudotime bins for all modalities was used to compute a dynamic time warping distance between all genes using the R package dtw^66^, which was further used for K means clustering with the R package FCPS^67^. For each of these clusters, we performed GO enrichment against the 6000 most highly variable genes from RNA. One of these clusters was highly enriched for neuron-related biological processes and the genes in this cluster were further used in subsequent analyses. Fig. 3a shows a subset of the clustering, that covers all pseudotemporal patterns. We provide the full list of genes in Supplementary Table 5.

### Reconstruction of the neurogenesis trajectory for the 4 months time point

To better understand pseudotemoral expression and chromatin modification dynamics during neurogenesis, we sought to further mitigate potential confounding factors such as distribution of time points, data quality and the pseudotime inference procedure. For this, we subset the data to telencephalic NPCs and neurons from the 4 month time point and applied the following additional quality filters to remove outliers: H3K27ac: > 1000 peak fragments per cell; H3K4me3: > 316 (10^2.5) peak fragments per cell, < 10000 peak fragments per cell. H3K27me3: < 3162 (10^3.5) peak fragments per cell. Pseudotime inference was subsequently performed using diffusion maps as described above. For RNA, we re-computed a set of 2000 variable features, excluding cell cycle genes, further regressed out cell cycle scores (ScaleData()) and performed PCA (RunPCA()). We used the first 20 PCs to compute diffusion maps with the R package destiny^30^. Analogous to the CUT&Tag data, ranks along the first diffusion component were used as a pseudotemoral ordering. The trajectory was then divided into 10 or 20 bins (depending on the analysis). Extended Data Fig. 7b shows a subset of the clustering, that covers the major pseudotemporal patterns. We provide the full list of genes in Supplementary Table 6.

### Detection of inflection point for neuronal genes

To examine at which point the CUT&Tag signal at neuronal genes (from clustering analysis described above) changes during neurogenesis relative to their expression, we sought to identify the first pseudotime bin in which the detection rate significantly diverges upwards (Fig. 3f). For this, we used the neurogenesis pseudotime from the 4 months time point divided into 10 bins and defined the first bin as the baseline. For each subsequent bin, we performed differential detection analysis against the baseline using a binomial linear model as described above. Multiple testing correction was performed using the Benjamini-Hochberg method. Based on this, we determined the first bin with FDR < 0.01 and log2 fold change > 1.5.

### Gene regulatory network analysis

To assess how the chromatin modifications shape the gene regulatory network (GRN) during neurogenesis we first inferred a GRN using Pando based on a dataset with integrated RNA and ATAC modalities over organoid development^14^. To select genomic regions for GRN inference, we performed region-to-gene linkage using the Signac function LinkPeaks() and intersected these regions with CUT&Tag peaks with >5% detection rate in any high-resolution cluster from all modalities. We used these regions as an input to the Pando function initiate_grn(). We further ran find_motifs() using the TF motifs provided by Pando and inferred a the GRN using infer_grn() with the following non-default parameters: peak_to_gene_method=’GREAT’, tf_cor=0.2, downstream=100000, and aggregated ATAC data to high-resolution clusters called by the authors (aggregate_peaks_col=’highres_clusters’). We filtered this network for significant positive connections with FDR > 0.01, pearson correlation > 0.2, coefficient > 0. To investigate the regulation of neurogenesis using this network, we extracted a subnetwork containing all neuronal genes (from clustering analysis described above) as well as TFs shared by at least three of these genes. For the regulatory regions in this network, we evaluated the epigenetic status in each pseudotime bin by applying a detection threshold of 5% for each chromatin modality. Network visualization and analysis was performed with ggraph (TLP; https://ggraph.data-imaginist.com/authors.html) and tidygraph (TLP; https://tidygraph.data-imaginist.com/authors.html).

### Analysis of multiome data of the developing human brain

We aligned the sequencing reads to the human genome and transcriptome (hg38, provided by 10x Genomics) using Cell Ranger Arc (version 2.0.0) with default parameters to obtain fragment files, peak calls and transcript counts. The data was further preprocessed and analyzed using Seurat (version 3.2)^56^ and Signac (version 1.1)^57^. Quality control was performed using the following criteria: 1000-6000 detected features per cell (RNA), >1500 UMI counts per cell (RNA), 1000-50000 detected peaks per cell (ATAC). Based on RNA expression, we further performed variable feature selection, z-scaling and PCA. The first 20 PCs were used as an input for Louvain clustering and UMAP. The clusters were manually annotated based on expression of marker genes. To specifically analyze the telencephalic neuron trajectory in this data, we extracted clusters corresponding to dorsal telencephalon NPCs, IPs and neurons. For this subset of the data, we again performed variable feature selection, z-scaling and PCA, and used the first 20 PCs to run UMAP and to compute diffusion maps with destiny^30^. Ranks along the first diffusion component were used as a pseudotemoral ordering. The velocity pseudotime was further rank-transformed and divided by the total number of cells in the dataset.

### Analysis of pseudotemporal and regional variance

To assess how expression and enrichment of genes and peaks varied during neurogenesis over pseudotime and between regions, we first selected 4000 highly variable genes and peaks with detection in >5% of cells in any high-resolution cluster. Next, we computed the average expression and activity for each high-resolution cluster. We fit three Gaussian linear models for each gene i module with mean cluster expression (Y) as the response variable and region assignment and/or pseudotime as the independent variables:

1. Y ∼ region
2. Y ∼ pseudotime
3. Y ∼ pseudotime + region

We used the value of these models as the fraction of variance explained by region (1), pseudotime (2) or branch and pseudotime (3).

### Distribution of chromatin states during differentiation pseudotime

To assess how genomic regions change chromatin states during differentiation, we first selected regions where peaks of the three marks were detected in >5% of cells in any pseudotime bin. For each mark, we further determined a detection threshold by computing the median detection for all peaks in these regions in all pseudotime bins. For each modality and chromatin bin we defined regions with peaks above this detection threshold as detected. If a region was marked by H3K27me3 and H3K4me3 in the same bin, they were annotated as bivalent and if they were marked by H3K27me3 and H3K27ac in the same bin, they were annotated as active promoters.

### Preprocessing and integration of drug treatment scRNA-seq data

The scRNA-seq data of A395 treated organoids was preprocessed analogous to the scRNA-seq data from the developmental time course. We then used Harmony^68^ with default parameters to integrate the different samples. Using the harmony integration, we performed Louvain clustering with a resolution of 0.2 and annotated the clusters based on canonical marker gene expression.

### Preprocessing of bulk CUT&Tag data

The fastq reads from the bulk CUT&Tag experiment were aligned to the human genome (hg38) using bwa^69^. Next, normalized bigwig files were obtained using deeptools bamCoverage^70^ with--ignoreDuplicates, -bs 200 and --normalizeUsing RPKM. To normalize bigwig files based on spike-ins, we first aligned spike-in reads to the human genome (hg38) using bwa^70^. Next, computed scaling factors from the resulting bam files using multiBamSummary bins with default parameters and used the result to perform normalization with bamCoverage^70^ (--ignoreDuplicates, -bs 200 and --scaleFactor). Heatmaps were generated using deeptools computeMatrix and plotHeatmap on the all H3K27me3 peaks from the stages PSC, NNE, NE of the developmental timecourse that were detected in >5% of the cells.

### Differential peak analysis in the perturbation experiment

To obtain enrichment scores on the level of peaks, we summarized normalized bigwig files to the peaks from the developmental time course for each modality. Based on these peak intensities, we computed the log_2_-fold change of treated versus control samples for each sample and concentration separately. The log_2_-fold changes were then summarized by computing the mean for each concentration.

### Differential gene expression analysis in the perturbation experiment

To assess the changes in gene expression upon treatment with A395, we performed differential expression (DE) analysis based using a logistic regression framework. To test for global DE while accounting for compositional differences, we considered the Louvain cluster label as a covariate in the model. We further accounted for sampling time point and sequencing depth (UMI count):

[Y_i sim n_UMI + time point + louvain_cluster + treatment]

We used the Seurat function FindMarkers() to perform the test for each condition separately and for all conditions combined. The resulting p-values were FDR-adjusted using the Benjamini-Hochberg method. We used a significance threshold of FDR < 0.01 and absolute log_2_-fold change > 0.1.

### Differential gene expression analysis

To test for compositional differences upon treatment with A395, we performed a Cochran-Mantel-Haenzel test stratified by sampling time point for each Louvain cluster and concentration separately. The resulting p-value was FDR-corrected and a significance threshold of 10^-4^ was applied.

### Calculation of repression, activation and bivalency scores

To assess the epigenetic state of DEG in different cell states, we computed repression (H3K27me3) activation (H3K4me3) or bivalency (H3K27me3 & H3K4me3) across high-resolution clusters. To compute activation and repression scores we used the Seurat function FindModuleScore() to compute the gene activity deviation from background for H3K4me3 and H3K27me3. We computed the means of these scores for each high resolution cluster and subsequently z-scaled them. A bivalency score was defined as zscale(H3K4me3 score) - zscale(H3K27me3 score). Here, a value close to 0 indicates bivalency, while positive and negative values indicate predominant activation (H3K4me3) and repression (H3K27me3), respectively.

### Inference of terminal fate probabilities in the perturbation experiment

To better resolve the differentiation hierarchies in the perturbation data, we computed transition probabilities into terminal fates based on RNA velocity. Count matrices for the spliced and unspliced transcriptome were obtained using kallisto (version 0.46.0)^63^ by running the command line tool loompy fromfastq from the python package loompy (version 3.0.6)(https://linnarssonlab.org/loompy/). Next, we used scVelo (version 0.2.4)^22^ and scanpy (version 1.8.2)^64^ to perform RNA velocity analysis. The 2000 most highly variable features were selected using the function scanpy.pp.highly_variable_genes() and used to compute moments in PCA space using the function scvelo.pp.moments() with n_neighbors=20. RNA velocity was calculated using the function scvelo.tl.velocity() with mode=’stochastic’ and a velocity graph was constructed using scvelo.tl.velocity_graph() with default parameters. To order cells in the developmental trajectory, a root cell was chosen randomly from cells of the first time point (EB) and velocity pseudotime was computed with scvelo.tl.velocity_pseudotime(). We next computed fate probabilities into the following manually annotated terminal cell states: Non-neural ectoderm, neural crest and neurons. For this, we used CellRank (version 1.3.0)^41^. A transition matrix was constructed with a palantir kernel (PalantirKernel()) based on velocity pseudotime. Absorption probabilities for each of the predefined terminal states were computed using the GPCCA estimator. Fate probabilities for each cell were visualized using a circular projection^71^. In brief, we evenly spaced terminal states around a circle and assigned each state an angle α_*t*_. We then computed 2D coordinates (*x*_*i*_, *y*_*i*_) from the *F ∈ R^Nxnt^* transition probability matrix for N cells and *n*_*t*_ terminal states as

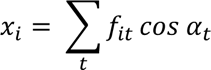

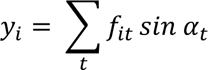

To visualize enrichment of perturbed cells in this space, we used the method outlined in Nikolova et al.^72^. Here, the kNN graph (k=100) was computed using euclidean distances in fate probability space and enrichment scores were visualized on the circular projection. Otherwise, the method was performed as described in the preprint.

### Transcription factor motif enrichment

To find transcription factors with putative involvement in aberrant cell fate determination upon EED inhibitor treatment, we first determined positions of binding motifs in peak regions. Position weight matrices (PWM) of human TF binding motifs were obtained from the CORE collection of JASPAR2020^73^. Motif positions in peak regions were determined using the R package motifmatchr (version 1.14)^74^ through the Signac function FindMotifs(). Next, we selected genomic regions in proximity (<10kb distance) to genes with differential expression in ectopic clusters (FDR < 10-4, log2 fold change > 1) that were depleted upon treatment (log2 fold change < −1). We used a Fisher exact test to obtain p-values for differential enrichment of motifs in these regions. The p-values were FDR corrected and a significance threshold of 0.05 was applied.

## Acknowledgments

We thank all members of the Camp and Treutlein laboratories, A. Jain and R. Holtackers for advice on imaging, Z. He for general bioinformatics support and R. Okamoto and M. Seimiya for general cell culture support, the lab of N. Iovino for initial help and discussion in establishing the protocol, in particular D. Ibarra Morales, N. Atinbayeva, F. Cardamone and E. Loeser. Y. Zhan for discussions. S. Riesenberg and S. Pääbo for providing the iCRISPR cell lines; staff at the Institute for Ophthalmology Basel for providing the 01F49i-N-B7 cell line. Illumina sequencing was performed by I. Nissen, E. Vogel Burcklen and C. Beisel at the Genomics Facility at D-BSSE, ETH Zurich. We are grateful to the Microscope facility at D-BSSE, ETH and E. Montani and T. Lummen for training. This work was supported by Chan Zuckerberg Initiative DAF, an advised fund of the Silicon Valley Community Foundation CZF2019-002440 (to J.G.C. and B.T.), the European Research Council (803441-Anthropoid, to J.G.C.; 758877-Organomics, to B.T.; 874606-Braintime, to B.T.), the Swiss National Science Foundation (project grant 310030_84795, to J.G.C.; project grant 310030_192604, to B.T.), and the Swiss National Center of Competence in Research Molecular Systems Engineering (to B.T.). J.S.F. was supported by the Boehringer Ingelheim Fonds. F.Z. was supported by EMBO Long-Term Fellowship ALTF 36-2021.

## Author contributions

F.Z. generated organoids used in this study, with support from B.K. and M.S.. M.S. helped with iPS cell culture. F.Z generated all single-cell transcriptome and single-cell CUT&Tag datasets with support from S.J. and performed initial bioinformatics analysis of all data. F.Z. performed the drug-treatment, bulk CUT&Tag and Western Blot experiments. F.Z. performed immunofluorescence experiments with support from B.E.. J.S.F. performed the analysis of the scRNA-seq/CUT&Tag-seq developmental time course with support from F.Z. J.D. helped in obtaining primary human brain samples and in the preparation of single cell suspensions. J.S.F. analyzed the drug treatment scRNA-seq and and bulk CUT&Tag data with support from F.Z.. F.Z., B.T. and J.G.C. designed the study and F.Z., J.S.F., B.T. and J.G.C. wrote the manuscript.

## Materials & Correspondence

All processed data are available at https://episcape.ethz.ch All code generated in the study including analysis parameters is available at https://github.com/quadbiolab/organoid_epigenomics. All experimental materials are available upon request to

---

**Extended Data Fig. 1.**
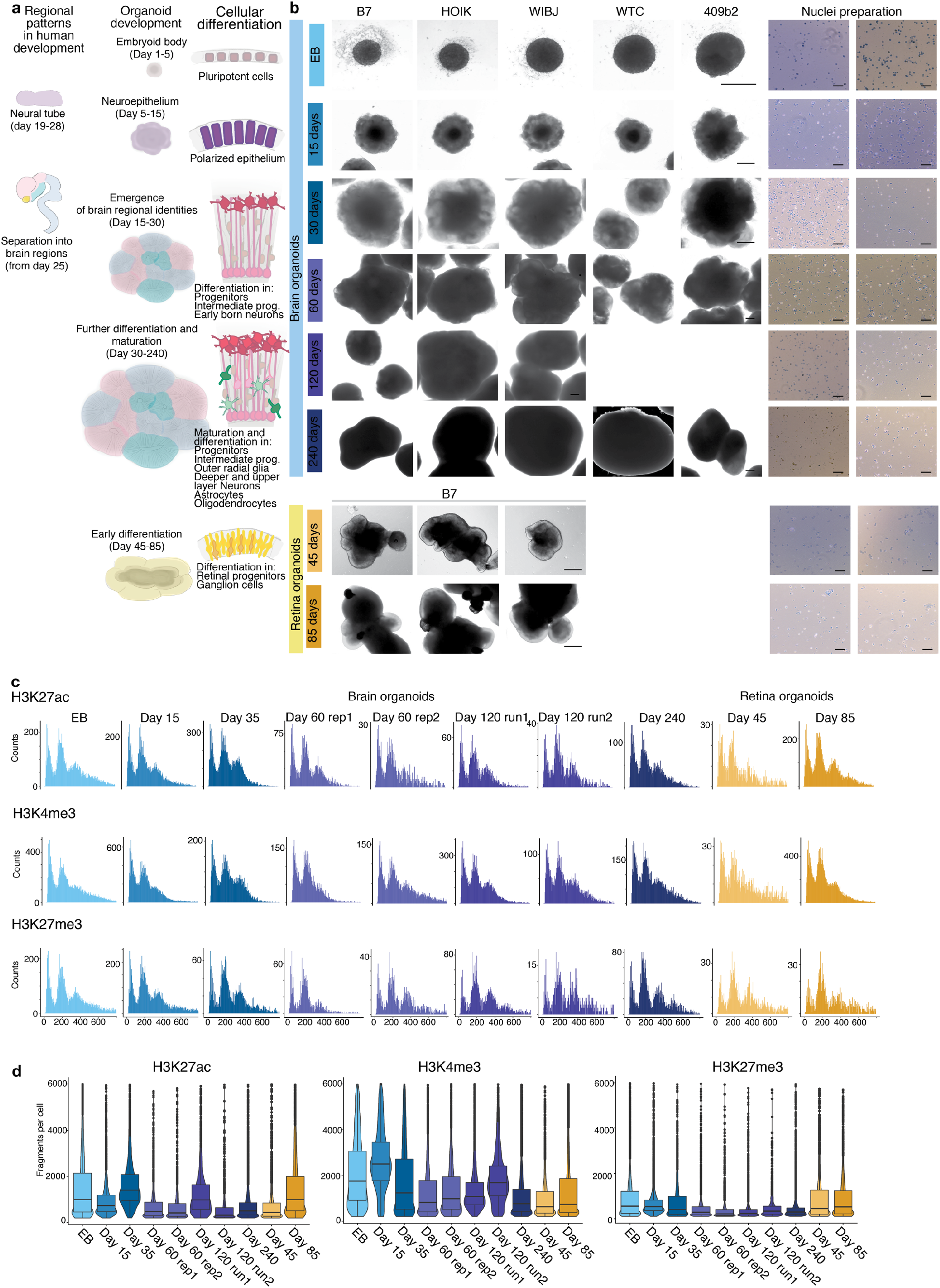
Epigenetic profiling of brain organoid development. a, Schematic of brain and retina organoid development in relation to early human brain development and emergence of cell type heterogeneity. b, Brightfield images of organoids during development (scale bar 500 µm) and the corresponding nuclei suspension (scale bar 100 µm). c, Histograms showing fragment length distribution for all samples and modalities. d, Violin plots showing the distribution of detected fragments per cell for each time point and chromatin modality.

**Extended Data Fig. 2.**
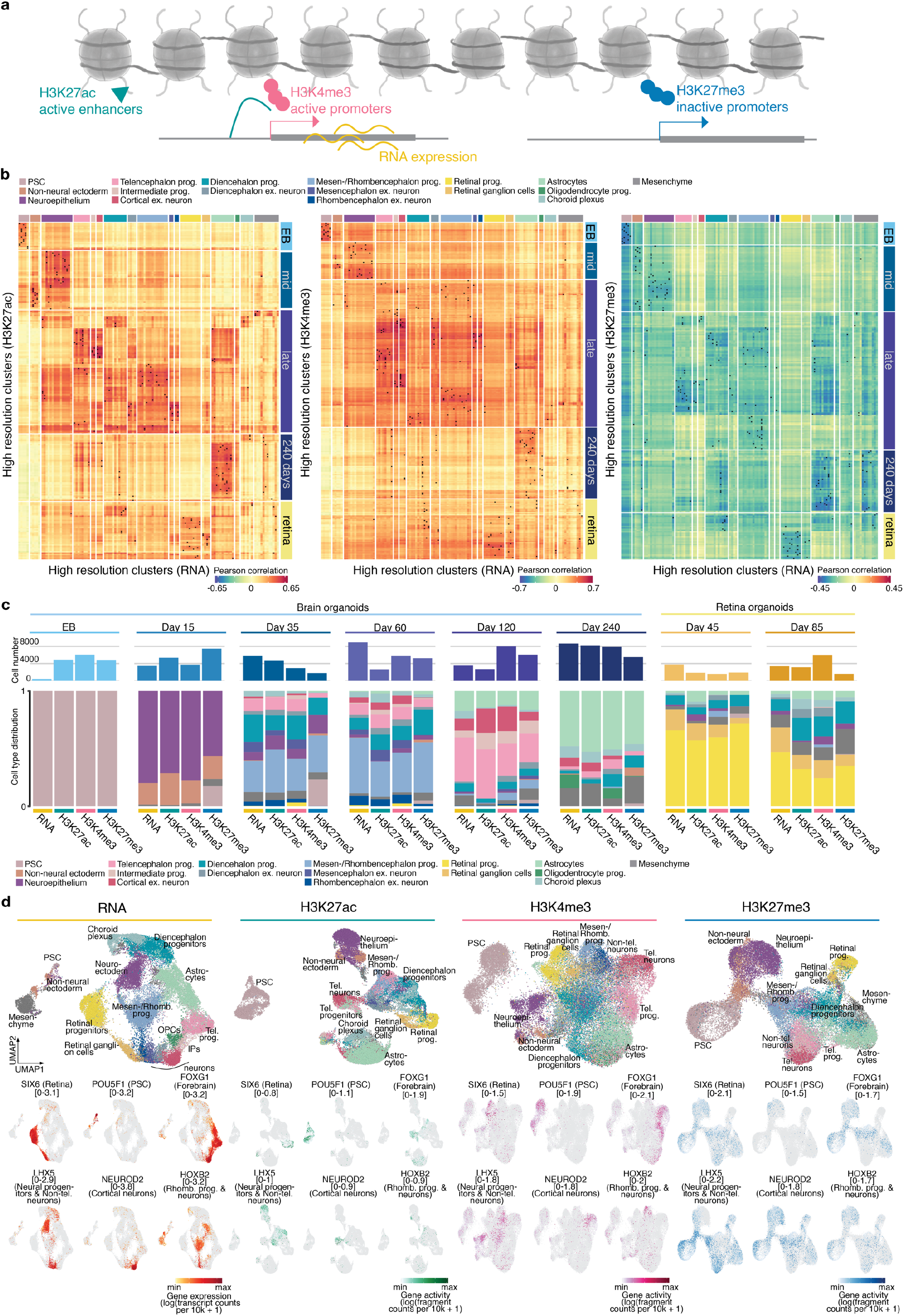
Multi-modal matching between high-resolution clusters. a, Schematic of the profiled modalities and their molecular role. H3K27ac marks active promoters and enhancers, H3K4me3 marks active promoters. H3K27me3 marks repressed regions with inactive promoters. b, Heatmaps showing correlation and matching of high-resolution clusters across modalities. Matched clusters are indicated with a cross. c, Bar-plot of absolute cell number per time point (top panel) for all modalities. Stacked bar plot of cell type distribution within each time point (bottom panel) (ex. - excitatory) for all modalities. d, UMAP representation of all modalities in the developmental time course colored by cell state (top panel) expression (bottom panel) (log(transcript counts per 10k + 1); in case of RNA-seq) and gene activity (log(fragment counts per 10k + 1); in case of CUT&Tag of genes marking different regional identities and cell states (Non-tel. – Non-telencephalon)

**Extended Data Fig. 3.**
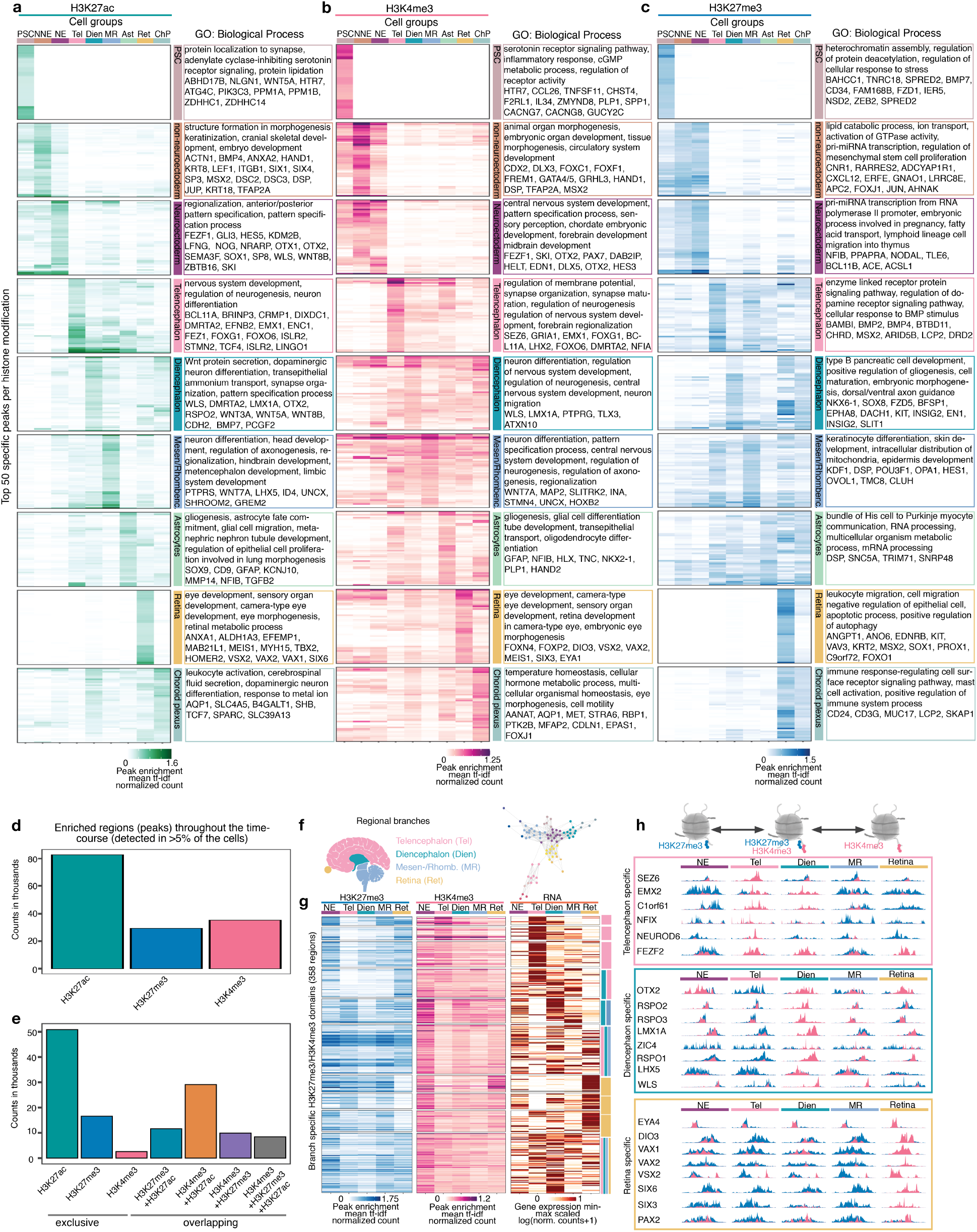
Cell type- and region-specific chromatin modification peaks. a-c Heatmaps showing normalized signal enrichment at state- and region-specific peaks for H3K27ac (a), H3K4me3 (b), and H3K27me3 (c). Selected genes in proximity to these peaks and representative GO Biological Process terms are indicated on the right. Tf-idf, term frequency-inverse document frequency. d, Barplot of the total number of peaks detected in more than 5% of the cells for each group of epigenetic modifications over all time points. e, Barplot of the number of peaks detected in more than 5% of the cells exclusive for each group of epigenetic modifications and overlapping between modifications over all time points. f, Schematic of the human brain (left) and force directed-layout of the main regional branches (right). g, Heatmap of peak enrichment of lineage-specific peaks showing switching between H3K27me3 and H3K4me3 marks. Expression of the closest gene is shown in the right panel. h, Genomic tracks showing bivalent peaks close to lineage-specific genes in the neuroepithelium that switch to activation or repression in the regional branches (blue-H3K27me3, magenta-H3K4me3).

**Extended Data Fig. 4.**
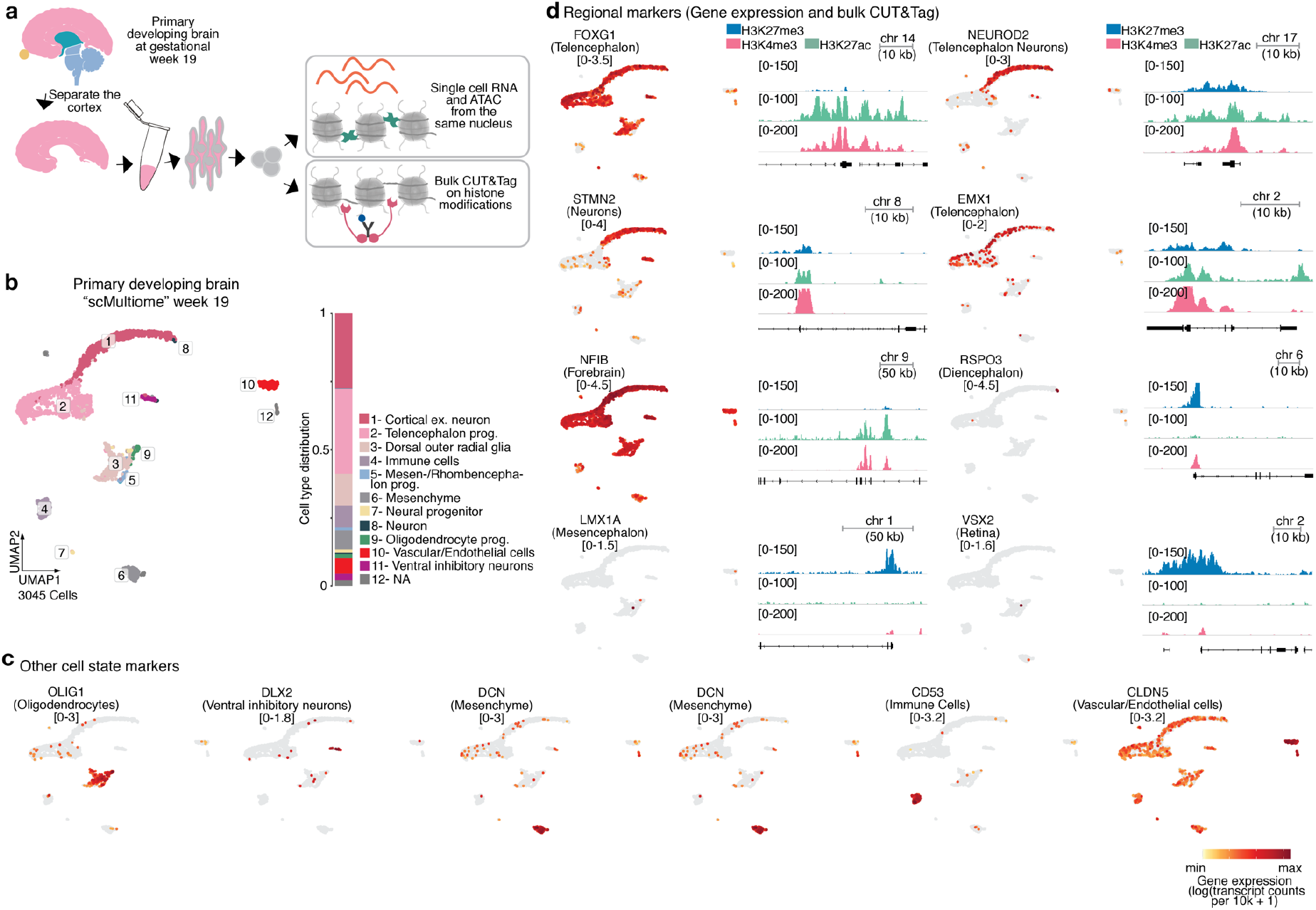
Single-cell “multiome” and bulk CUT&Tag data of human developing cortex at gestation week 19. a, Schematic of the experimental setup. b, UMAP embedding colored by cell state of the “scMultiome” data from the human developing brain (right). Barplot showing the cell state distribution within the sample reveals that most cells originate from the telencephalon. c, UMAP embedding as in (b) colored by gene expression of cell state markers used to annotate the different cell populations. d, Regional markers identified in Fig. 2e-f recapitulate the switching of activating and repressing marks in signal tracks of bulk H3K27me3, H3K27ac and H3K4me3 CUT&Tag data from a primary developing telencephalon (FOXG1, NEUROD2, STMN2, EMX1, NFIB) while non-telencephalon regional markers remain repressed (RSPO3, LMX1A, VSX2).

**Extended Data Fig. 5.**
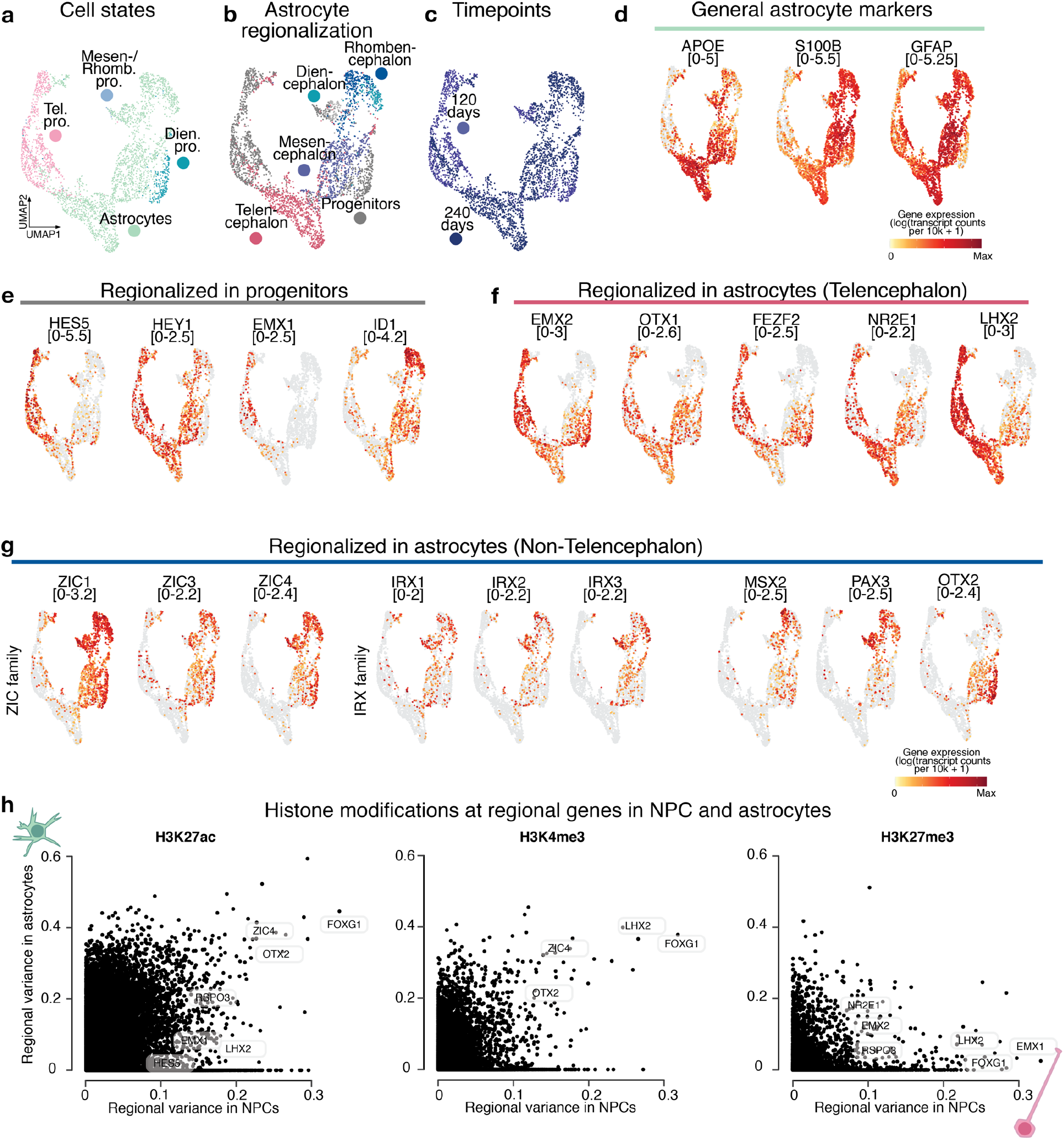
Astrocyte regionalization signatures in human brain organoids. a-c, UMAP embedding of NPCs and astrocytes from the 120 and 240 day time points colored by cell state (a), brain region (b), or time point (c). d, UMAP embedding colored by gene expression of general astrocyte marker genes. e-g, UMAP embedding colored by expression of genes that show a brain region specific expression pattern in NPCs (e), are specifically expressed in forebrain astrocytes and NPCs (f) are specifically expressed in non-forebrain astrocytes and NPCs (g). h, Scatter plot comparing the regional variance of histone modification enrichment (for H3K27ac, H3K4me3, H3K27me3) between astrocytes and NPCs.

**Extended Data Fig. 6.**
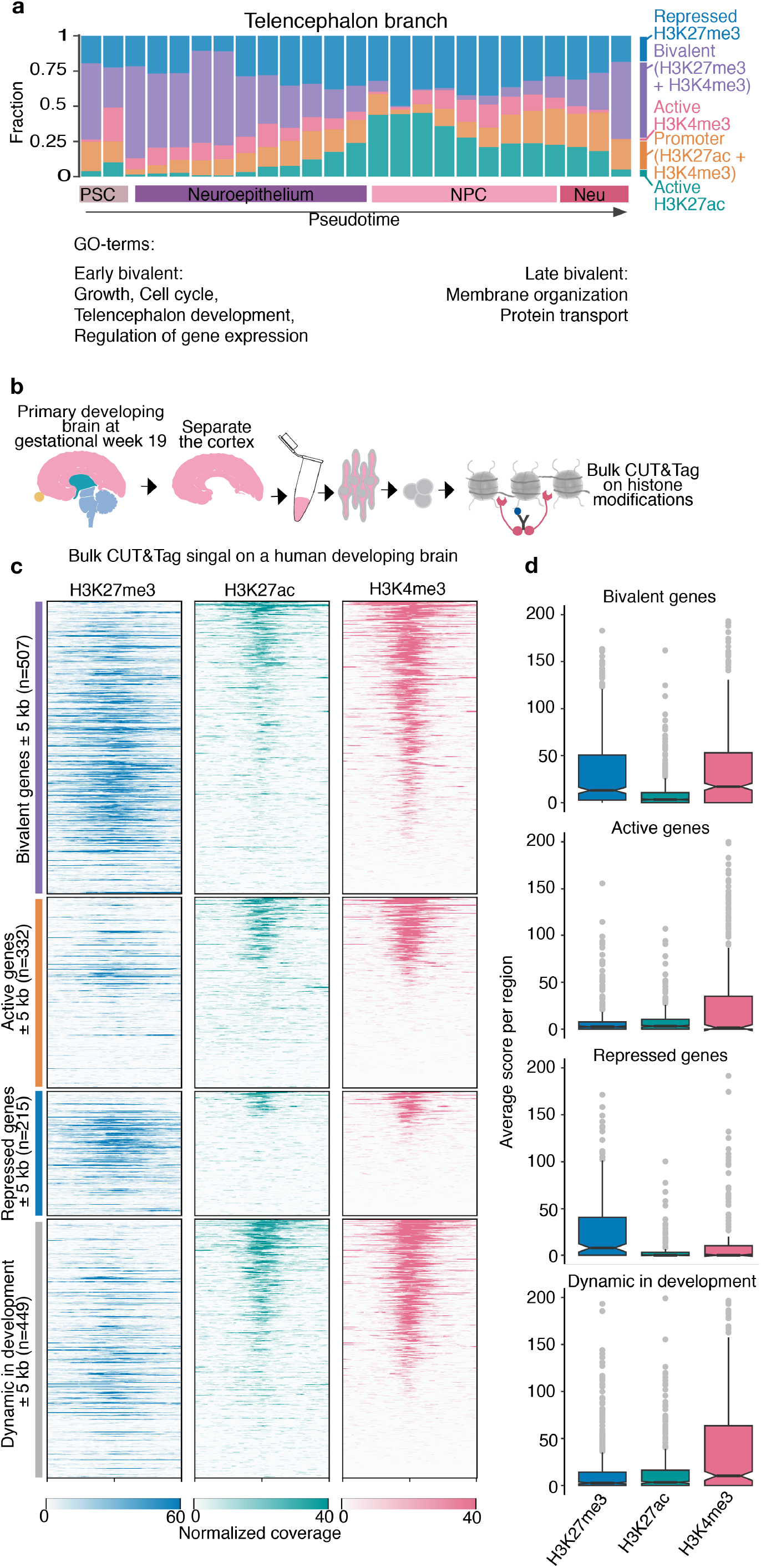
Chromatin states of the telencephalon branch in organoid and primary data. a, Stacked barplot showing the distribution of different chromatin states over pseudotime from PSC to neurons (see Materials and Methods for details.) Bivalent domains are enriched in PSCs and become again abundant at the neuronal stage. GO term analysis of genes close to bivalent domains reveals enrichment for growth, cell cycle, telencephalon development, regulation of gene expression at early stages (PSC and neuroepithelium) and membrane organization and protein transport at later stages (NPC and neuron). b, Schematic of the experiment. c, H3K27me3, H3K27ac and H3K4me3 bulk CUT&Tag signal from a primary developing cortex at gw 19 plotted on the chromatin states from (a), (bivalent, active, repressed and dynamically changing). Regions from (a) were summarized starting with NPC to neurons. We plotted the signal on the TSS of the gene closest to the domain. For the categories active and repressed, only genes that were exclusively active/repressed in 90% of bins were considered. d, quantification of the signal of the respective epigenetic marks in the corresponding cluster.

**Extended Data Fig. 7.**
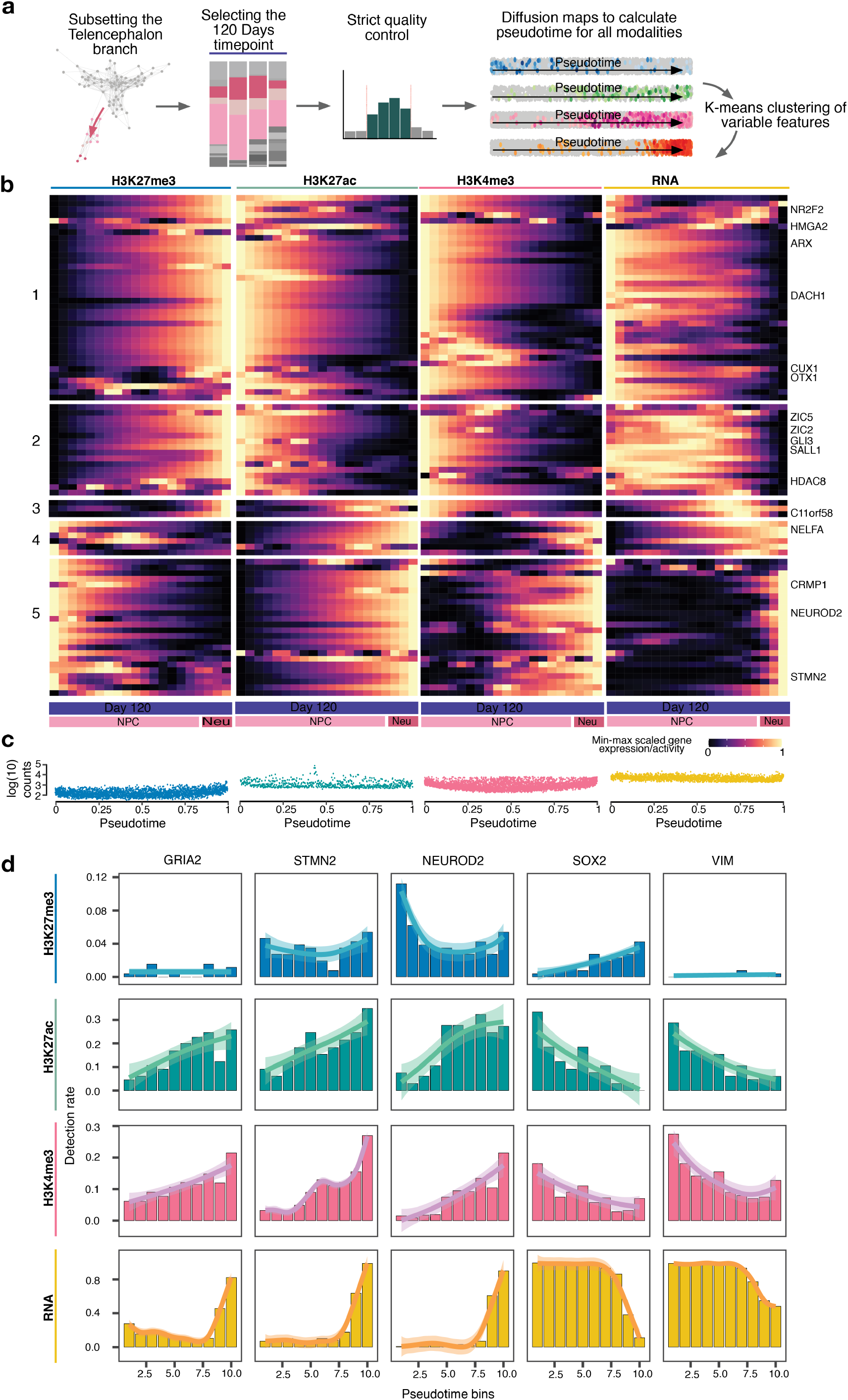
Pseudotemporal reconstruction of dorsal telencephalic neuron differentiation using diffusion maps. a, Schematic of the analysis. We subset the telencephalon branch and isolated only cells of the 120 days time point. The cells were again filtered and pseudotime for all modalities was calculated using diffusion maps. b, Heatmap (left) showing gene activity scores for H3K27me3, H3K27ac and H3K4me3 as well as RNA expression over the telencephalic neuron differentiation trajectory from pluripotency. Pseudotime was binned and genes were K-means clustered based on average expression/activity of all marks and RNA in all bins (Supplementary Table 6 contains all clusters). Using this strictly refined analysis we could again identify a neuronal cluster that clearly showed priming with active chromatin modifications. c, Jitter plot showing even UMI and fragment count distribution over pseudotime for all modalities. d, Barplot of detection rate over pseudotime at selected example genes.

**Extended Data Fig. 8.**
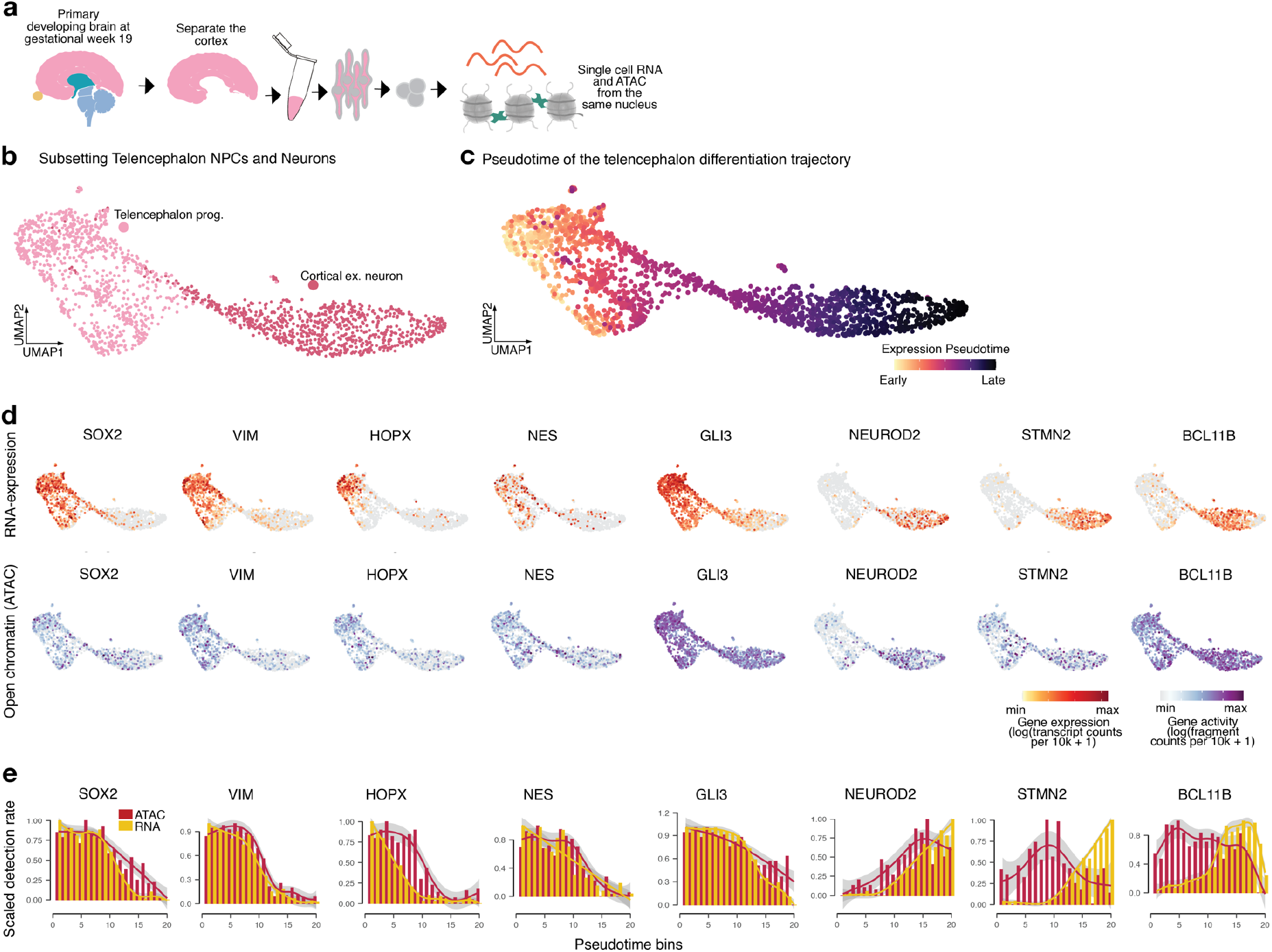
Pseudotime reconstruction shows dynamic RNA expression and chromatin accessibility changes during neurogenesis in the primary developing cortex. a, Schematic of the experiment. b, UMAP embedding colored by cell state showing the telencephalic NPCs and neurons. c, UMAP embedding colored by pseudotime. d, UMAP embedding as is (b) coloured by expression of different neuronal and progenitor marker genes (top) and chromatin openness (bottom). e, Detection rate of scATAC signal and scRNA expression measured from the same cells of a primary human developing embryonic brain at gestation week 19. Neuronal genes (NEUROD2, STMN2, BCL11B) show priming of chromatin.

**Extended Data Fig. 9.**
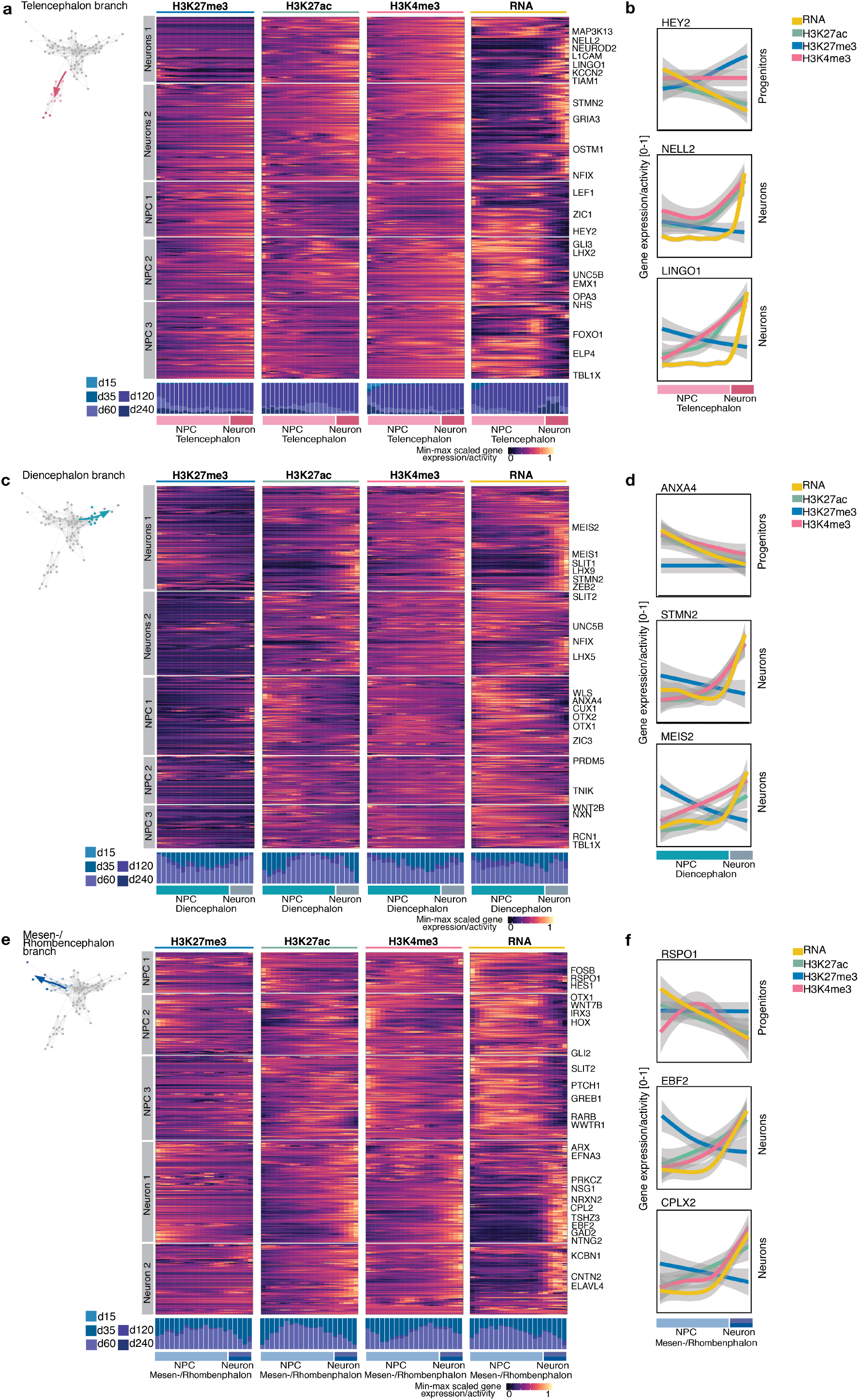
Pseudotime reconstruction of dynamic changes of all modalities during neurogenesis in different developing organoid brain regions. a, Heatmap showing gene activity scores for H3K27me3, H3K27ac and H3K4me3 as well as RNA expression over the telencephalic neuron differentiation trajectory from NPCs. Pseudotime was binned and genes were K-means clustered based on average expression/activity of all marks and RNA in all bins. (The averaged pseudotemporal gene expression and activities of cluster 1 are shown as line plots in Fig. 3j, as well as GO term enrichment) b, Line plots showing smoothed pseudotemporal expression and activities of selected examples from multiple K-means clusters. HEY2 is expressed at the progenitor state, showing alignment of pseudotimes. NELL2 and LINGO1 are expressed in neurons and exhibit epigenetic priming with activating marks. c, Same as (a) for the diencephalic neuron differentiation trajectory from NPCs. (The averaged pseudotemporal gene expression and activities of cluster 1 are shown as line plots in Fig. 3j, as well as GO term enrichment) d, Line plots showing smoothed pseudotemporal expression and activities of selected examples from multiple K-means clusters. ANXA4 is expressed at the progenitor state, showing alignment of pseudotimes. STMN2 and MEIS2 are expressed in neurons and exhibit epigenetic priming with activating marks. e, Same as (a) for the rhomben- and mesencephalic neuron differentiation trajectory from NPCs. (The averaged pseudotemporal gene expression and activities of cluster 4 are shown as line plots in Fig. 3j, as well as GO term enrichment) f, Line plots showing smoothed pseudotemporal expression and activities of selected examples from multiple K-means clusters. RSPO1 is expressed at the progenitor state, showing alignment of pseudotimes. EBF2 and CPLX2 are expressed in neurons and exhibit epigenetic priming with activating marks.

**Extended Data Fig. 10.**
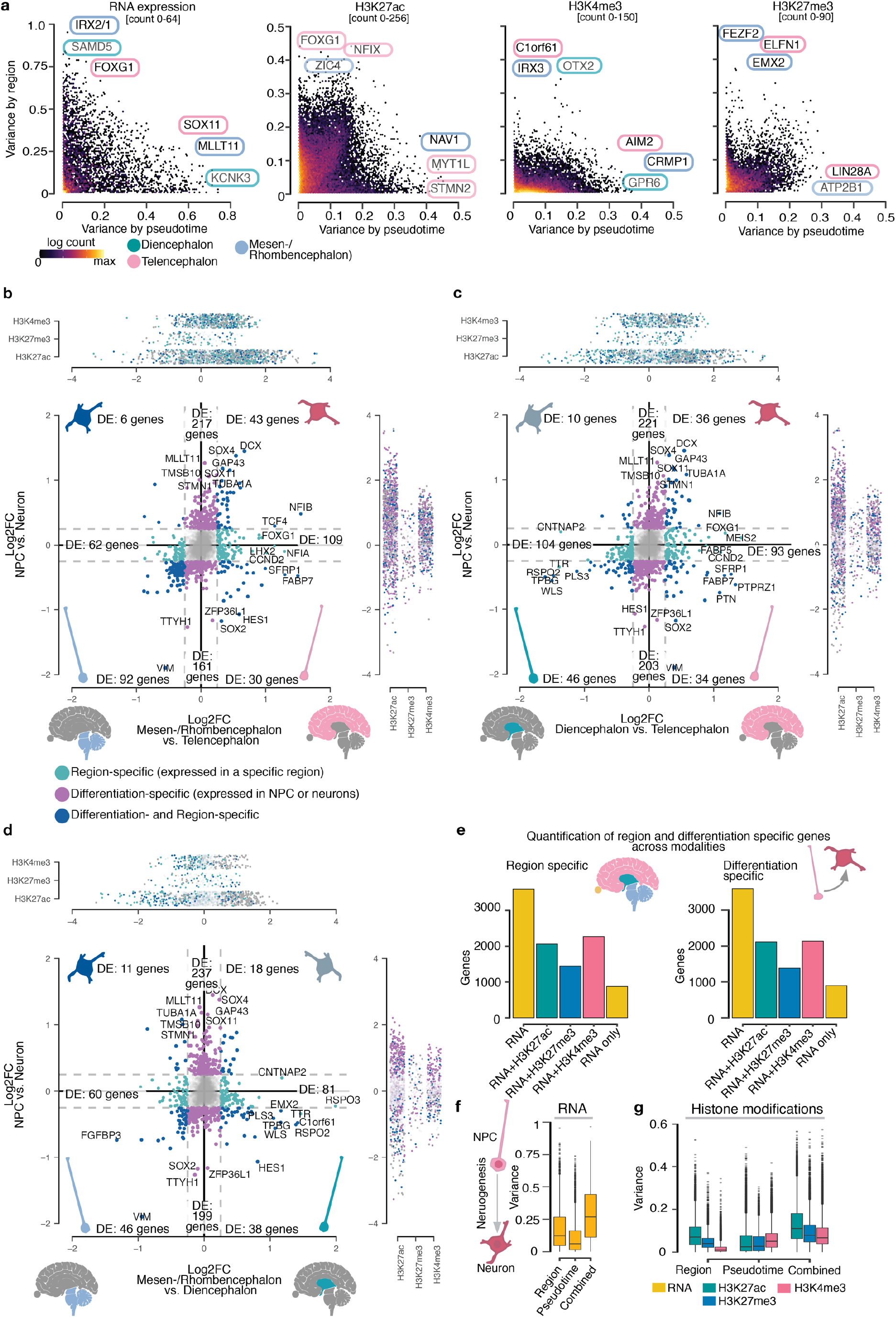
Comparing differentiation- and region-specific gene expression between different brain regional branches. a, Scatter plot of the per gene/peak expression variance by pseudotime and per gene/peak expression by the brain regional branch for RNA (top) and all histone modifications. b, Scatter plot showing the log_2_FC of gene expression between neurons and progenitors (y-axis) and telencephalon and mesen/rhombencephalon (x-axis). The scatter-plots on the right and top show the same gene set comparing the log_2_FC of the individual histone modifications. c, Scatter plot showing the log_2_FC of gene expression between neurons and progenitors (y-axis) and diencephalon and telencephalon (x-axis). The scatter-plots on the right and top show the same gene set comparing the log_2_FC of the individual histone modifications. d, Scatter plot showing the log_2_FC of gene expression between neurons and progenitors (y-axis) and mesen/rhombencephalon and diencephalon (x-axis). The scatter-plots on the right and top show the same gene set comparing the log_2_FC of the individual histone modifications. e, Barplot showing the overlap for region and differentiation specific genes detected with each modality (RNA and histone modifications). f, Boxplot of the gene expression variance for the brain regional branch, pseudotime and combined. g, Boxplot of variance of the histone modification enrichment per peak for the brain regional branch, pseudotime and combined.

**Extended Data Fig. 11.**
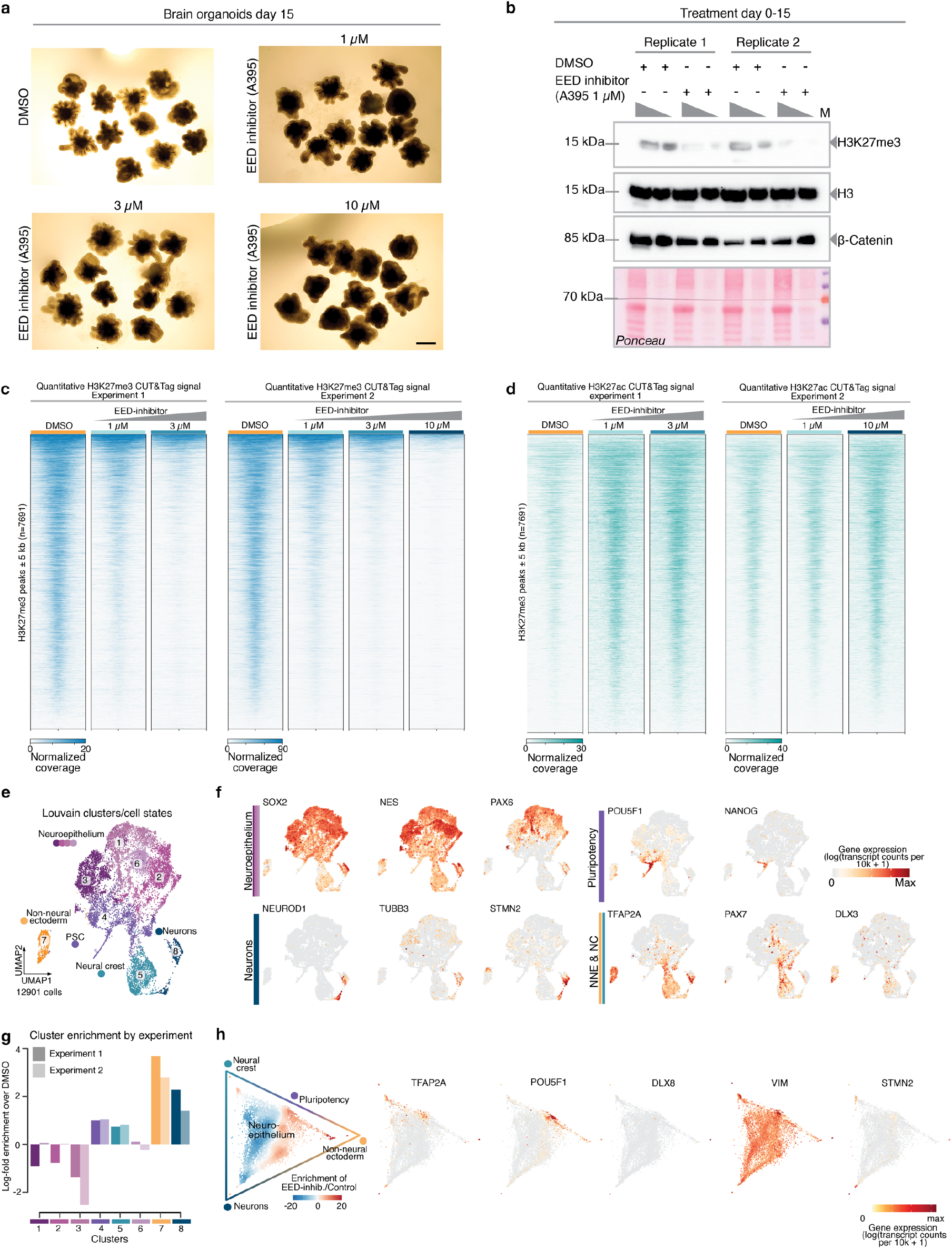
Aberrant cell fate acquisition upon depletion of H3K27me3. a, Organoids at day 15 treated with DMSO (control) or different concentrations of A395 an EED inhibitor. b, Western Blot of cellular extracts of 15 day old organoids shows depletion of H3K27me3 upon treatment with 1 µM A395. H3, β-Catenin and Ponceau serve as loading controls. c, Heatmap of bulk H3K27me3 CUT&Tag signal on H3K27me3 peaks of two replicates of the inhibitor treatment, showing consistent depletion of H3K27me3. d, Heatmap of bulk H3K27ac CUT&Tag signal on H3K27me3 peaks of two replicates of the inhibitor treatment, showing consistent increase of H3K27ac. e, UMAP embedding colored by cluster annotation of the DMSO control and all inhibitor treated cells. f, UMAP embedding colored by expression of genes marking annotated populations from (e). g, Barplot showing the two replicates of the EED inhibitor treatment. h, Circular plot of differential transition probabilities between different cell states from the inhibitor treatment, showing an enrichment of H3K27me3 depleted cells at the terminal states of the graph (left), the circular plot colored by the expression of different marker genes (right).

**Extended Data Fig. 12.**
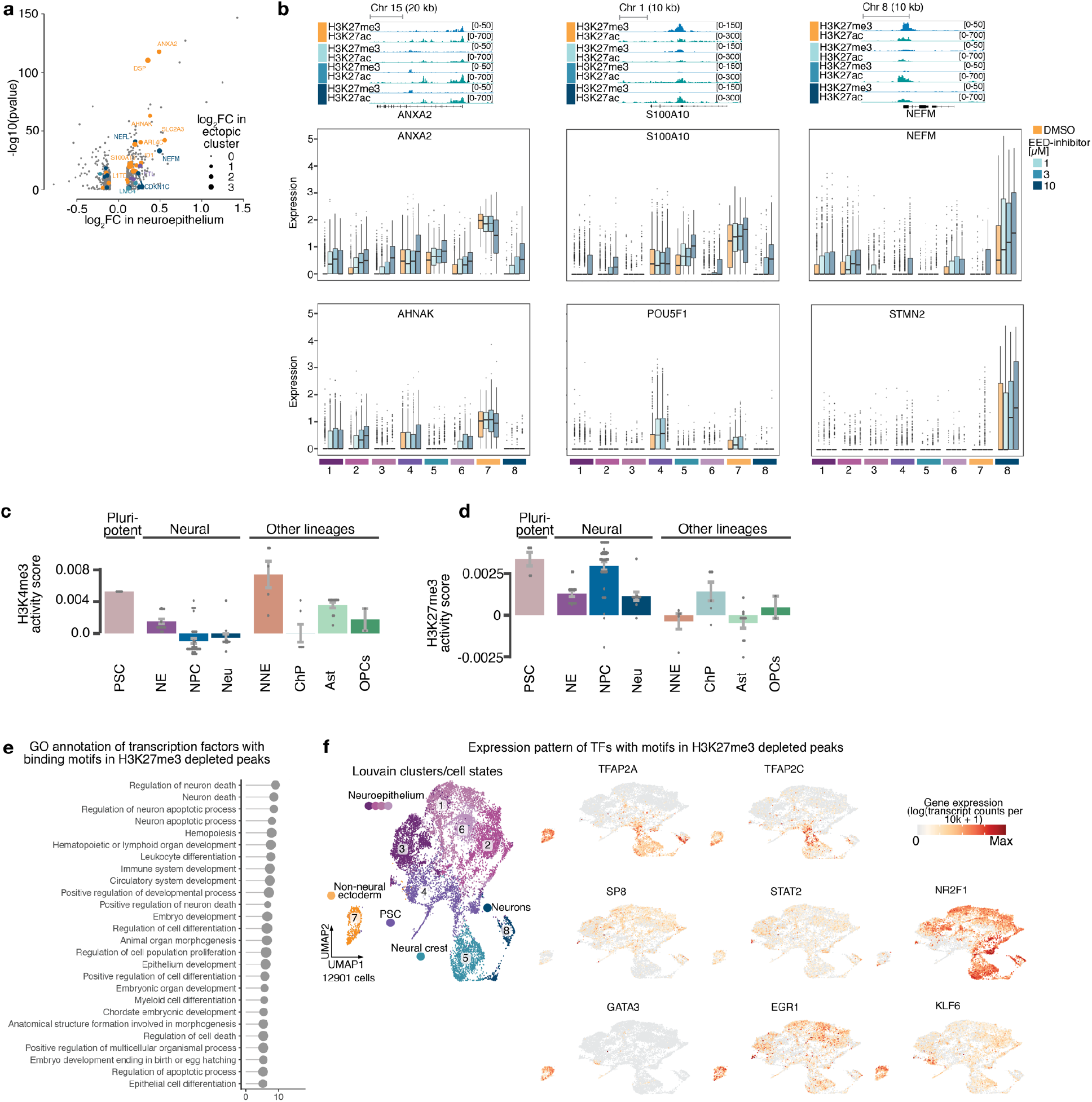
Effect of H3K27me3 depletion on lineage commitment in human brain organoids. a, Vulcano plot of the log2FC between the inhibitor treatment and the controls in the neuroepithelium clusters and the –log10(pvalue). Genes are colored by the identity of the ectopically formed clusters where they act as cluster markers. b, IGV browser snapshot showing the inhibitor concentration dependent reduction of H3K27me3 and gain of H3K27ac at selected targets from (a) (top panel) Boxplot of upregulated genes in the neural epithelium showing gene expression per cluster (bottom panel). c, Barplots quantifying the activity score of H3K4me3 on the top 200 DE genes between inhibitor treatment and and control. d, same as c for H3K27me3. e, GO term enrichment of transcription factors corresponding to binding motifs enriched in H3K27me3 depleted peaks in proximity to DE genes. f, UMAP embedding colored by cluster annotation of the DMSO control and all inhibitor treated cells (left panel). UMAP embedding colored by gene expression for several transcription factors with enriched binding motifs (Fig. 4j) in H3K27me3 depleted peaks close to deregulated genes (right panel).

## References and Notes

1 Millan-Zambrano, G., Burton, A., Bannister, A. J. & Schneider, R. Histone post-translational modifications - cause and consequence of genome function. Nat Rev Genet 23, 563–580, doi:10.1038/s41576-022-00468-7 (2022).

2 Paulsen, B., et al. Autism genes converge on asynchronous development of shared neuron classes. Nature 602, 268–273, doi:10.1038/s41586-021-04358-6 (2022).

3. Li, C. et al. Single-cell brain organoid screening identifies developmental defects in autism. bioRxiv, 2022.2009.2015.508118, doi:10.1101/2022.09.15.508118 (2022).

4 Preissl, S., Gaulton, K. J. & Ren, B. Characterizing cis-regulatory elements using single-cell epigenomics. Nat Rev Genet, doi:10.1038/s41576-022-00509-1 (2022).

5 Schuettengruber, B., Bourbon, H. M., Di Croce, L. & Cavalli, G. Genome Regulation by Polycomb and Trithorax: 70 Years and Counting. Cell 171, 34–57, doi:10.1016/j.cell.2017.08.002 (2017).

6 Rada-Iglesias, A., et al. A unique chromatin signature uncovers early developmental enhancers in humans. Nature 470, 279–283, doi:10.1038/nature09692 (2011).

7 Creyghton, M. P., et al. Histone H3K27ac separates active from poised enhancers and predicts developmental state. Proc Natl Acad Sci U S A 107, 21931–21936, doi:10.1073/pnas.1016071107 (2010).

8 Bernstein, B. E., et al. A bivalent chromatin structure marks key developmental genes in embryonic stem cells. Cell 125, 315–326, doi:10.1016/j.cell.2006.02.041 (2006).

9 Voigt, P., Tee, W. W. & Reinberg, D. A double take on bivalent promoters. Genes Dev 27, 1318–1338, doi:10.1101/gad.219626.113 (2013).

10 Zenk, F., et al. Germ line-inherited H3K27me3 restricts enhancer function during maternal-to-zygotic transition. Science 357, 212–216, doi:10.1126/science.aam5339 (2017).

11 Lavarone, E., Barbieri, C. M. & Pasini, D. Dissecting the role of H3K27 acetylation and methylation in PRC2 mediated control of cellular identity. Nature Communications 10, 1679, doi:10.1038/s41467-019-09624-w (2019).

12 Becht, E., et al. Dimensionality reduction for visualizing single-cell data using UMAP. Nat Biotechnol, doi:10.1038/nbt.4314 (2018).

13 Kanton, S., et al. Organoid single-cell genomic atlas uncovers human-specific features of brain development. Nature 574, 418–422, doi:10.1038/s41586-019-1654-9 (2019).

14 Fleck, J. S., et al. Inferring and perturbing cell fate regulomes in human brain organoids. Nature, doi:10.1038/s41586-022-05279-8 (2022).

15 Kelava, I., Chiaradia, I., Pellegrini, L., Kalinka, A. T. & Lancaster, M. A. Androgens increase excitatory neurogenic potential in human brain organoids. Nature 602, 112–116, doi:10.1038/s41586-021-04330-4 (2022).

16 Rowitch, D. H. & Kriegstein, A. R. Developmental genetics of vertebrate glial-cell specification. Nature 468, 214–222, doi:10.1038/nature09611 (2010).

17 Bartosovic, M., Kabbe, M. & Castelo-Branco, G. Single-cell CUT&Tag profiles histone modifications and transcription factors in complex tissues. Nat Biotechnol 39, 825–835, doi:10.1038/s41587-021-00869-9 (2021).

18 Wu, S. J. et al. Single-cell CUT&Tag analysis of chromatin modifications in differentiation and tumor progression. Nat Biotechnol 39, 819–824, doi:10.1038/s41587-021-00865-z (2021).

19 Zhu, C., et al. Joint profiling of histone modifications and transcriptome in single cells from mouse brain. Nat Methods 18, 283–292, doi:10.1038/s41592-021-01060-3 (2021).

20 Blondel, V. D., Guillaume, J.-L., Lambiotte, R. & Lefebvre, E. Fast unfolding of communities in large networks. Journal of Statistical Mechanics: Theory and Experiment 2008, P10008, doi:10.1088/1742-5468/2008/10/p10008 (2008).

21 Stark, S. G., et al. SCIM: universal single-cell matching with unpaired feature sets. Bioinformatics 36, i919–i927, doi:10.1093/bioinformatics/btaa843 (2020).

22 Bergen, V., Lange, M., Peidli, S., Wolf, F. A. & Theis, F. J. Generalizing RNA velocity to transient cell states through dynamical modeling. Nat Biotechnol 38, 1408–1414, doi:10.1038/s41587-020-0591-3 (2020).

23 McLean, C. Y., et al. GREAT improves functional interpretation of cis-regulatory regions. Nat Biotechnol 28, 495–501, doi:10.1038/nbt.1630 (2010).

24 Jadhav, U., et al. Acquired Tissue-Specific Promoter Bivalency Is a Basis for PRC2 Necessity in Adult Cells. Cell 165, 1389–1400, 10.1016/j.cell.2016.04.031 (2016).

25 Welle, A., et al. Epigenetic control of region-specific transcriptional programs in mouse cerebellar and cortical astrocytes. Glia 69, 2160–2177, doi:10.1002/glia.24016 (2021).

26 Herrero-Navarro, Á., et al. Astrocytes and neurons share region-specific transcriptional signatures that confer regional identity to neuronal reprogramming. Sci Adv 7, doi:10.1126/sciadv.abe8978 (2021).

27 Zeisel, A., et al. Molecular Architecture of the Mouse Nervous System. Cell 174, 999–1014.e1022, doi:10.1016/j.cell.2018.06.021 (2018).

28. Braun, E. et al. Comprehensive cell atlas of the first-trimester developing human brain. bioRxiv, 2022.2010.2024.513487, doi:10.1101/2022.10.24.513487 (2022).

29 Batiuk, M. Y., et al. Identification of region-specific astrocyte subtypes at single cell resolution. Nat Commun 11, 1220, doi:10.1038/s41467-019-14198-8 (2020).

30 Haghverdi, L., Buttner, M., Wolf, F. A., Buettner, F. & Theis, F. J. Diffusion pseudotime robustly reconstructs lineage branching. Nat Methods 13, 845–848, doi:10.1038/nmeth.3971 (2016).

31 Burns, A. M. & Gräff, J. Cognitive epigenetic priming: leveraging histone acetylation for memory amelioration. Current Opinion in Neurobiology 67, 75–84, 10.1016/j.conb.2020.08.011 (2021).

32 Ziffra, R. S., et al. Single-cell epigenomics reveals mechanisms of human cortical development. Nature 598, 205–213, doi:10.1038/s41586-021-03209-8 (2021).

33 Ma, S., et al. Chromatin Potential Identified by Shared Single-Cell Profiling of RNA and Chromatin. Cell 183, 1103–1116.e1120, doi:10.1016/j.cell.2020.09.056 (2020).

34 Lara-Astiaso, D. et al. Immunogenetics. Chromatin state dynamics during blood formation. Science 345, 943–949, doi:10.1126/science.1256271 (2014).

35 Zeller, P., et al. Single-cell sortChIC identifies hierarchical chromatin dynamics during hematopoiesis. Nat Genet 55, 333–345, doi:10.1038/s41588-022-01260-3 (2023).

36 Wang, H., et al. H3K4me3 regulates RNA polymerase II promoter-proximal pause-release. Nature 615, 339–348, doi:10.1038/s41586-023-05780-8 (2023).

37 He, Y., et al. The EED protein-protein interaction inhibitor A-395 inactivates the PRC2 complex. Nat Chem Biol 13, 389–395, doi:10.1038/nchembio.2306 (2017).

38 Sankar, A., et al. Histone editing elucidates the functional roles of H3K27 methylation and acetylation in mammals. Nat Genet 54, 754–760, doi:10.1038/s41588-022-01091-2 (2022).

39 Loh, C. H., van Genesen, S., Perino, M., Bark, M. R. & Veenstra, G. J. C. Loss of PRC2 subunits primes lineage choice during exit of pluripotency. Nat Commun 12, 6985, doi:10.1038/s41467-021-27314-4 (2021).

40 Pereira, J. D., et al. Ezh2, the histone methyltransferase of PRC2, regulates the balance between self-renewal and differentiation in the cerebral cortex. Proc Natl Acad Sci U S A 107, 15957–15962, doi:10.1073/pnas.1002530107 (2010).

41 Lange, M., et al. CellRank for directed single-cell fate mapping. Nature Methods 19, 159–170, doi:10.1038/s41592-021-01346-6 (2022).

42 Hoffman, T. L., Javier, A. L., Campeau, S. A., Knight, R. D. & Schilling, T. F. Tfap2 transcription factors in zebrafish neural crest development and ectodermal evolution. J Exp Zool B Mol Dev Evol 308, 679–691, doi:10.1002/jez.b.21189 (2007).

43 Williams, T., Admon, A., Luscher, B. & Tjian, R. Cloning and expression of AP-2, a cell-type-specific transcription factor that activates inducible enhancer elements. Genes Dev 2, 1557–1569, doi:10.1101/gad.2.12a.1557 (1988).

44 Ciceri, G. et al. An epigenetic barrier sets the timing of human neuronal maturation. bioRxiv, 2022.2006.2002.490114, doi:10.1101/2022.06.02.490114 (2022).

45 Argelaguet, R., et al. Multi-omics profiling of mouse gastrulation at single-cell resolution. Nature 576, 487–491, doi:10.1038/s41586-019-1825-8 (2019).

46 Muñoz-Sanjuán, I. & Brivanlou, A. H. Neural induction, the default model and embryonic stem cells. Nat Rev Neurosci 3, 271–280, doi:10.1038/nrn786 (2002).

47 Riesenberg, S., et al. Simultaneous precise editing of multiple genes in human cells. Nucleic Acids Res 47, e116, doi:10.1093/nar/gkz669 (2019).

48 Kilpinen, H., et al. Common genetic variation drives molecular heterogeneity in human iPSCs. Nature 546, 370–375, doi:10.1038/nature22403 (2017).

49 Roberts, B., et al. Systematic gene tagging using CRISPR/Cas9 in human stem cells to illuminate cell organization. Molecular Biology of the Cell 28, 2854–2874, doi:10.1091/mbc.e17-03-0209 (2017).

50 Cowan, C. S., et al. Cell Types of the Human Retina and Its Organoids at Single-Cell Resolution. Cell 182, 1623–1640 e1634, doi:10.1016/j.cell.2020.08.013 (2020).

51 Giandomenico, S. L., Sutcliffe, M. & Lancaster, M. A. Generation and long-term culture of advanced cerebral organoids for studying later stages of neural development. Nat Protoc 16, 579–602, doi:10.1038/s41596-020-00433-w (2021).

52 Kaya-Okur, H. S. et al. CUT&Tag for efficient epigenomic profiling of small samples and single cells. Nat Commun 10, 1930, doi:10.1038/s41467-019-09982-5 (2019).

53 Picelli, S., et al. Tn5 transposase and tagmentation procedures for massively scaled sequencing projects. Genome Res 24, 2033–2040, doi:10.1101/gr.177881.114 (2014).

54 Buenrostro, J. D., Wu, B., Chang, H. Y. & Greenleaf, W. J. ATAC-seq: A Method for Assaying Chromatin Accessibility Genome-Wide. Curr Protoc Mol Biol 109, 21 29 21–21 29 29, doi:10.1002/0471142727.mb2129s109 (2015).

55 Stoeckius, M., et al. Simultaneous epitope and transcriptome measurement in single cells. Nat Methods 14, 865–868, doi:10.1038/nmeth.4380 (2017).

56 Stuart, T., et al. Comprehensive Integration of Single-Cell Data. Cell 177, 1888–1902 e1821, doi:10.1016/j.cell.2019.05.031 (2019).

57 Stuart, T., Srivastava, A., Madad, S., Lareau, C. A. & Satija, R. Single-cell chromatin state analysis with Signac. Nat Methods 18, 1333–1341, doi:10.1038/s41592-021-01282-5 (2021).

58 Kang, H. M., et al. Multiplexed droplet single-cell RNA-sequencing using natural genetic variation. Nat Biotechnol 36, 89–94, doi:10.1038/nbt.4042 (2018).

59. Wahle, P. et al. Multimodal spatiotemporal phenotyping of human organoid development. bioRxiv, 2022.2003.2016.484396, doi:10.1101/2022.03.16.484396 (2022).

60 He, Z., Brazovskaja, A., Ebert, S., Camp, J. G. & Treutlein, B. CSS: cluster similarity spectrum integration of single-cell genomics data. Genome Biology 21, 224, doi:10.1186/s13059-020-02147-4 (2020).

61 Fleck, J. S., et al. Resolving organoid brain region identities by mapping single-cell genomic data to reference atlases. Cell Stem Cell 28, 1148–1159.e1148, doi:10.1016/j.stem.2021.02.015 (2021).

62 Virtanen, P., et al. SciPy 1.0: fundamental algorithms for scientific computing in Python. Nature Methods 17, 261–272, doi:10.1038/s41592-019-0686-2 (2020).

63 Bray, N. L., Pimentel, H., Melsted, P. & Pachter, L. Near-optimal probabilistic RNA-seq quantification. Nature Biotechnology 34, 525–527, doi:10.1038/nbt.3519 (2016).

64 Wolf, F. A., Angerer, P. & Theis, F. J. SCANPY: large-scale single-cell gene expression data analysis. Genome Biology 19, 15, doi:10.1186/s13059-017-1382-0 (2018).

65 Wood, S. N. Fast stable restricted maximum likelihood and marginal likelihood estimation of semiparametric generalized linear models. Journal of the Royal Statistical Society: Series B (Statistical Methodology) 73, 3–36 (2011).

66 Giorgino, T. Computing and Visualizing Dynamic Time Warping Alignments in R: The dtw Package. Journal of Statistical Software 31, 1–24, doi:10.18637/jss.v031.i07 (2009).

67 Thrun, M. C. & Stier, Q. Fundamental clustering algorithms suite. SoftwareX 13, 100642, 10.1016/j.softx.2020.100642 (2021).

68 Korsunsky, I., et al. Fast, sensitive and accurate integration of single-cell data with Harmony. Nature Methods 16, 1289–1296, doi:10.1038/s41592-019-0619-0 (2019).

69 Li, H. & Durbin, R. Fast and accurate short read alignment with Burrows-Wheeler transform. Bioinformatics 25, 1754–1760, doi:10.1093/bioinformatics/btp324 (2009).

70 Ramírez, F., et al. deepTools2: a next generation web server for deep-sequencing data analysis. Nucleic Acids Res 44, W160–165, doi:10.1093/nar/gkw257 (2016).

71 Velten, L., et al. Human haematopoietic stem cell lineage commitment is a continuous process. Nat Cell Biol 19, 271–281, doi:10.1038/ncb3493 (2017).

72 Nikolova, M. T. et al. Fate and state transitions during human blood vessel organoid development. bioRxiv, 2022.2003.2023.485329, doi:10.1101/2022.03.23.485329 (2022).

73 Fornes, O. et al. JASPAR 2020: update of the open-access database of transcription factor binding profiles. Nucleic Acids Research 48, D87–D92, doi:10.1093/nar/gkz1001 (2019).

74 Schep, A. motifmatchr: fast motif matching in R. R package version 1 (2021).

